# Simultaneous detection of membrane contact dynamics and associated Ca^2+^ signals by reversible chemogenetic reporters

**DOI:** 10.1101/2023.12.28.573515

**Authors:** Paloma García Casas, Michela Rossini, Linnea Påvénius, Mezida Saeed, Nikita Arnst, Sonia Sonda, Matteo Bruzzone, Valeria Berno, Andrea Raimondi, Maria Livia Sassano, Luana Naia, Patrizia Agostinis, Mattia Sturlese, Barbara A. Niemeyer, Hjalmar Brismar, Maria Ankarcrona, Arnaud Gautier, Paola Pizzo, Riccardo Filadi

**Affiliations:** Department of Biomedical Sciences, University of Padua, Padua, Italy; Department of Biochemistry, Molecular Biology and Physiology, Faculty of Medicine, Unidad de Excelencia Instituto de Biología y Genética Molecular (IBGM), University of Valladolid and CSIC, Valladolid, Spain; Institute of Neuroscience, National Research Council (CNR), Padua, Italy; Science for Life Laboratory, Department of Women’s and Children’s Health, Karolinska Institutet, Stockholm, Sweden; Science for Life Laboratory, Department of Applied Physics, Royal Institute of Technology, Stockholm, Sweden; ALEMBIC, IRCCS San Raffaele Scientific Institute, Milan, Italy; Cell Death Research and Therapy lab, Department of Cellular and Molecular Medicine, and Center for Cancer Biology-VIB, KU, Leuven, Belgium; Department of Neurobiology, Care Sciences and Society, Division of Neurogeriatrics, Center for Alzheimer Research, Karolinska Institutet, Stockholm, Sweden; Department of Pharmaceutical Science, University of Padua, Padua, Italy; Molecular Biophysics, Saarland University, Homburg, Germany; Sorbonne Université, École Normale Supérieure, Université PSL, CNRS, Laboratoire des Biomolécules, LBM, 75005 Paris, France; Centro Studi per la Neurodegenerazione (CESNE), University of Padua, Padua, Italy

**Author notes:** These authors contributed equally: Paloma García Casas, Michela Rossini.

## Abstract

Membrane contact sites (MCSs) are hubs allowing various cell organelles to coordinate their activities. The dynamic nature of these sites and their small size hinder analysis by current imaging techniques. To overcome these limitations, we here designed a series of reversible chemogenetic reporters incorporating improved, low-affinity variants of splitFAST, and studied the dynamics of different MCSs at high spatiotemporal resolution, both *in vitro* and *in vivo*. We demonstrated that these versatile reporters suit different experimental setups well, allowing one to address challenging biological questions. Using these novel probes, we identified a hitherto unknown pathway, in which calcium (Ca^2+^) signalling dynamically regulates endoplasmic reticulum-mitochondria juxtaposition, characterizing the underlying mechanism. Finally, by integrating Ca^2+^-sensing capabilities into the splitFAST technology, we introduced PRINCESS (<u>PR</u>obe for <u>IN</u>terorganelle <u>C</u>a^2+^-<u>E</u>xchange <u>S</u>ites based on <u>S</u>plitFAST), an unprecedented class of reporters to simultaneously detect MCSs and measure the associated Ca^2+^ dynamics using a single biosensor.

---

In eukaryotes, membrane-bound organelles allow the separation of sometimes incompatible biochemical reactions. Nevertheless, cell organelles do not work as isolated structures but coordinate their activities by forming specialized platforms, called membrane contact sites (MCSs), whereby their membranes are closely juxtaposed (typically in the 10-30 nm range)^1,2^. At MCSs, key cell pathways take place, including the exchange of metabolites, signalling molecules and information^1^. The efficiency of these processes is indeed strongly favoured by the very close proximity between organelle membranes and is aided by protein-protein or protein-lipid interactions. Specific protein and lipid compositions characterize each type of MCS, ensuring the performance of their specialized functionalities^2^.

Remarkably, early disturbances in MCS structure and/or function have been described in different high-incidence disorders, such as neurodegenerative diseases and several cancers^3,4^. This suggests the tantalizing hypothesis that such alterations might underlie pathogenesis, prompting an increasing interest in studying MCSs and the associated metabolic/signalling activities.

However, a comprehensive study of MCSs is hampered by the limitations of the currently available techniques. Indeed, because of the dynamic nature of MCSs^1,2^, their study requires monitoring changes that occur in response to specific needs with high spatiotemporal resolution, something that current tools do not provide^2,5^. For instance, optical microscopy allows to follow gross changes over time, but its diffraction-limited spatial resolution does not match the nanometric size of most MCSs. Conversely, electron microscopy (EM) and super-resolution microscopy can resolve membrane proximity in space, but need specialized equipment, are limited to fixed samples (*e.g*., EM) and/or require long acquisitions, which are poorly compatible with MCS dynamics. Recently, the introduction of self-complementing fluorescent probes based on split-GFP overcame the problem of spatial resolution and allowed the detection of MCSs^6,7^. However, in most of these systems, the complementation reaction leading to fluorescence maturation is slow (taking tens of minutes to hours) and poorly reversible^8^, possibly stabilizing MCSs and/or limiting the detection of MCS changes occurring within seconds. Moreover, current methods allow to study either MCS morphology (*i.e.*, the extent of inter-organelle apposition) or functionality (*i.e.*, the efficiency of a specific activity associated with MCSs, such as inter-organelle Ca^2+^ transfer), but no probe exists to simultaneously visualize MCSs with high spatial resolution and measure their activities over time.

To overcome these limitations, we generated a series of reversible chemogenetic reporters, focusing on the well characterized juxtaposition between endoplasmic reticulum (ER), mitochondria and the plasma membrane (PM). Specifically, we engineered splitFAST (split <u>F</u>luorescence-activating and <u>A</u>bsorption <u>S</u>hifting <u>T</u>ag), a small, split fluorescent protein tag recently introduced for the monitoring of dynamic protein-protein interactions^9^. The splitFAST fragments show modest affinity; however, when in close proximity, they assemble into a reporter that binds and stabilizes the fluorescent state of exogenously applied multicolour fluorogenic dyes (fluorogens) (**Fig. 1a**), which are otherwise dark when unbound. Remarkably, splitFAST complementation is rapid and promptly reversible^9^, making it an ideal system for studying MCS dynamics in physiological conditions.

**Figure 1:**
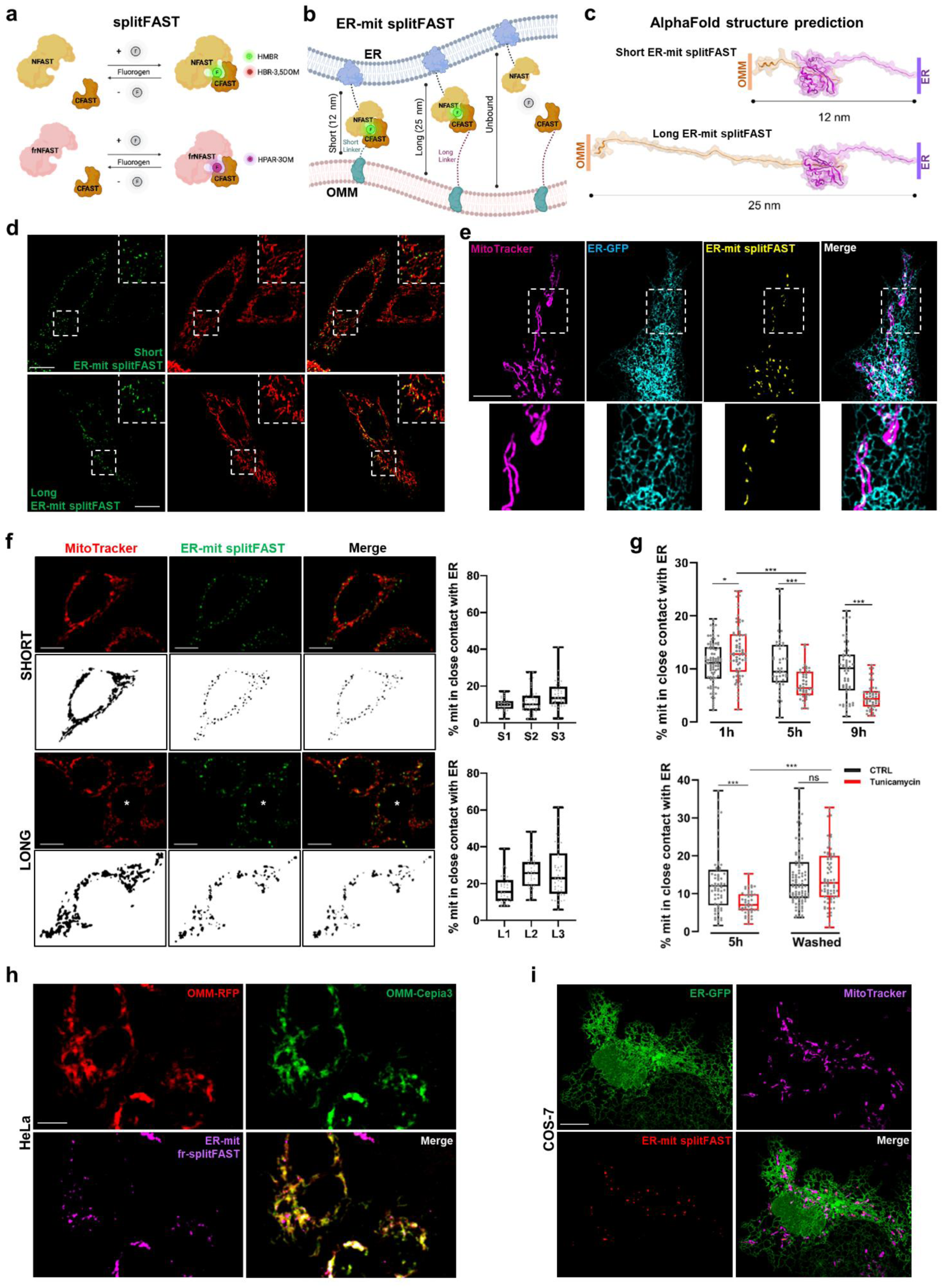
The splitFAST system for studying membrane contact site dynamics. **a**, The cartoon represents the splitFAST system, formed by the NFAST and the CFAST portions, that are not fluorescent *per se*, unless assembled and bound to a fluorogen: green, red or far-red (this latter requiring frNFAST to complement CFAST, see text). The fluorogens are dark when unbound, and the assembly of the complex is fully reversible. **b-c**, The cartoon (**b**) represents the tailoring of the splitFAST system to mark ER-mit MCSs. By targeting the NFAST portion to the ER membrane and CFAST10 to the OMM, both separated by unstructured linkers of different sizes as predicted by AlphaFold (**c**), two different probes have been generated to study ER-mit MCSs: the short ER-mit splitFAST, with a length of ∼12 nm, and the long ER-mit splitFAST, of ∼25 nm. **d**, Representative confocal images of HeLa cells co-expressing either the short or the long ER-mit splitFAST with a mit-RFP and exhibiting a dotted pattern, that colocalizes with the mitochondrial RFP-positive network. Scale bar, 10 μm. **e**, Representative confocal image of a COS-7 cell co-expressing the short ER-mit splitFAST and an ER-GFP, and stained for the mitochondrial network by MitoTracker Deep Red. A dotted ER-mit splitFAST signal colocalizes with both the ER and the mitochondria networks, marking sites of close contact between the two organelles. Scale bar, 10 μm. **f**, Representative confocal images of stable HeLa cell clones expressing either the short (S1, S2, S3) or the long (L1, L2, L3) version of the ER-mit splitFAST probes. A dotted signal along the mitochondrial network (labelled by MitoTracker Deep Red) is shown, allowing the calculation of the percentage of the mitochondrial surface (% mit) covered by either short or long contacts with the ER (box plots on the right, see Methods). The cell indicated with a white asterisk is shown in the corresponding binary image used for analysis. n=29-46 cells from 3 independent experiments. Scale bar, 10 μm. **g**, The box plots represent the dynamic changes of ER-mit MCSs (calculated as in panel **f**) in HeLa cells expressing the short ER-mit splitFAST, upon induction of ER stress with Tunicamycin (5 μg/ml). An increase in ER-mit MCSs was observed after 1 hour of treatment, followed by a decrease, after 5 and 9 hours. Recovery of basal levels of ER-mit MCSs was observed after the removal of the ER stressor. n>42 cells from 3 independent experiments.*p<0.05; **p<0.01; ***p<0.001. **h-i**, Representative confocal images of (**h)** HeLa cells co-expressing OMM-RFP, OMM-Cepia3 and short ER-mit fr-splitFAST probe (allowing to visualize ER-mit MCSs in far-red), or (**i**) of COS-7 cells co-expressing ER-GFP, short ER-mit splitFAST (allowing to visualize ER-mit MCSs in either red, this case, or green, see panel **d**) and in which mitochondrial network was marked with MitoTracker Deep Red. Scale bar, 10 μm.

We demonstrated that these reporters are endowed with high versatility, enabling to: a) detect MCSs *in vitro* and *in vivo*; b) follow complex MCS fusion/fission events associated with deep organelle and cytoskeleton remodelling; c) measure perturbations of organelle coupling in different physiological and pathological conditions; d) track the transient recruitment of proteins at MCSs; e) reveal hitherto unknown MCS rearrangements upon specific cell stimulations and characterize the underlying mechanisms.

Moreover, by tailoring the splitFAST system and combining it with suitable Ca^2+^-sensing modules, we generated a series of single reporters capable of simultaneously detecting MCSs and measuring the associated Ca^2+^ signals, that we dubbed PRINCESS (<u>PR</u>obe for <u>IN</u>terorganelle <u>C</u>a^2+^-<u>E</u>xchange <u>S</u>ites based on <u>S</u>plitFAST). By integrating unprecedented spatiotemporal resolution, fluorescence imaging and Ca^2+^-sensing capabilities, we showed that PRINCESS allows the study of MCS morphology and function using a single probe, opening new avenues for designing innovative biosensors enabling to address fundamental biological questions.

## Results

### Design of splitFAST-based probes to detect MCSs *in vitro* and *in vivo*

SplitFAST is a fluorescence complementation system formed by two parts, NFAST and CFAST, previously designed through bisection of the 14-kDa protein FAST for real-time visualization of transient protein-protein interactions^9,10^ (**Fig. 1a**). CFAST fragments of respectively 10 (CFAST10) or 11 (CFAST11) amino acids have been reported, with the former displaying a lower self-complementation and thus a higher dynamic range^9^. We firstly confirmed the prompt reversibility of NFAST-CFAST10 interaction (**Supplementary Fig. 1a** and **Supplementary Video 1**). Then, to test whether splitFAST can be tailored to detect MCSs, while preserving its dynamicity, we targeted NFAST and CFAST10 to the ER- and the outer mitochondrial membrane (OMM), respectively (**Fig. 1b**). Disordered linkers of different lengths were inserted between the OMM targeting sequence and CFAST10, generating two probes to detect either short (short ER-mit splitFAST) or long (long ER-mit splitFAST) ER-mitochondria contacts, potentially complementing at distances between ER and OMM of, respectively, less than 12 nm and 25 nm, as predicted by Alphafold^11^ (**Fig. 1c**). The expression of these probes in HeLa (**Fig. 1d**) and COS-7 (**Fig. 1e**) cells revealed a punctate pattern overlapping with the mitochondrial network, in correspondence of regions of ER-mitochondria co-localization (**Fig. 1e**). The fluorescent dots appeared few seconds after the addition of the fluorogen (**Supplementary Fig. 1b**), confirming the high cell-permeability of these molecules and the prompt splitFAST complementation. Different HeLa cell clones, expressing either the short or the long ER-mit splitFAST probe, displayed a larger fraction of mitochondrial surface engaged in the formation of long contacts with the ER, compared to the short ones (**Fig. 1f and Supplementary Fig. 1c**). Moreover, the treatment of cells expressing the short probe with the ER-stressor tunicamycin induced a transient increase, followed by a decrease, of ER-mitochondria contacts (ER-mit MCSs; **Fig. 1g**), in line with previous data^12^ and confirming that the splitFAST-based probes can dynamically follow MCS changes. Consistently, either decreased or increased ER-mit MCSs were associated with different cell treatments/conditions (**Supplementary Fig. 1d**). SplitFAST complementation at ER-mit MCSs was visualized by three different hydroxybenzylidene rhodanine (HBR)-derived fluorogens: HMBR (green fluorescence; **Fig. 1d**), HPAR-3OM (far-red fluorescence; **Fig. 1h**) and HBR-3,5DOM (red fluorescence, **Fig. 1i**), with HPAR-3OM requiring the replacement of NFAST with frNFAST (far-red NFAST^13^; see Methods and **Fig. 1a**). Notably, the high spectral flexibility of the system enables the combined study of MCSs with multiple fluorescent reporters and represents an advantage compared with previous techniques.

Importantly, an ideal method to study MCSs should not interfere with their dynamics. The choice of CFAST10, rather than CFAST11, is instrumental to this purpose, as it is endowed with a very low affinity for NFAST^9^. To further minimize possible MCS conditioning, we introduced in our probes RspA-splitFAST, an improved, orthologous splitFAST version endowed with even lower self-complementation, greater dynamic range and higher brightness^14^. RspA-NFAST and RspA-CFAST fragments were targeted, respectively, to the ER membrane and the OMM (hereafter referred as ER-mit RspA-splitFAST), giving rise to a bright, dotted fluorescent signal (**Fig. 2a**), precisely at regions corresponding to close ER-mitochondria juxtaposition, as revealed by correlative light electron microscopy (CLEM) experiments (**Fig. 2b and Supplementary Video 2**). Importantly, by this approach, we could verify that the fraction of mitochondrial perimeter in close contact with the ER was similar in control (untransfected) and ER-mit RspA-splitFAST-expressing cells (**Fig. 2b**), indicating that the probe does not force MCS formation.

**Figure 2:**
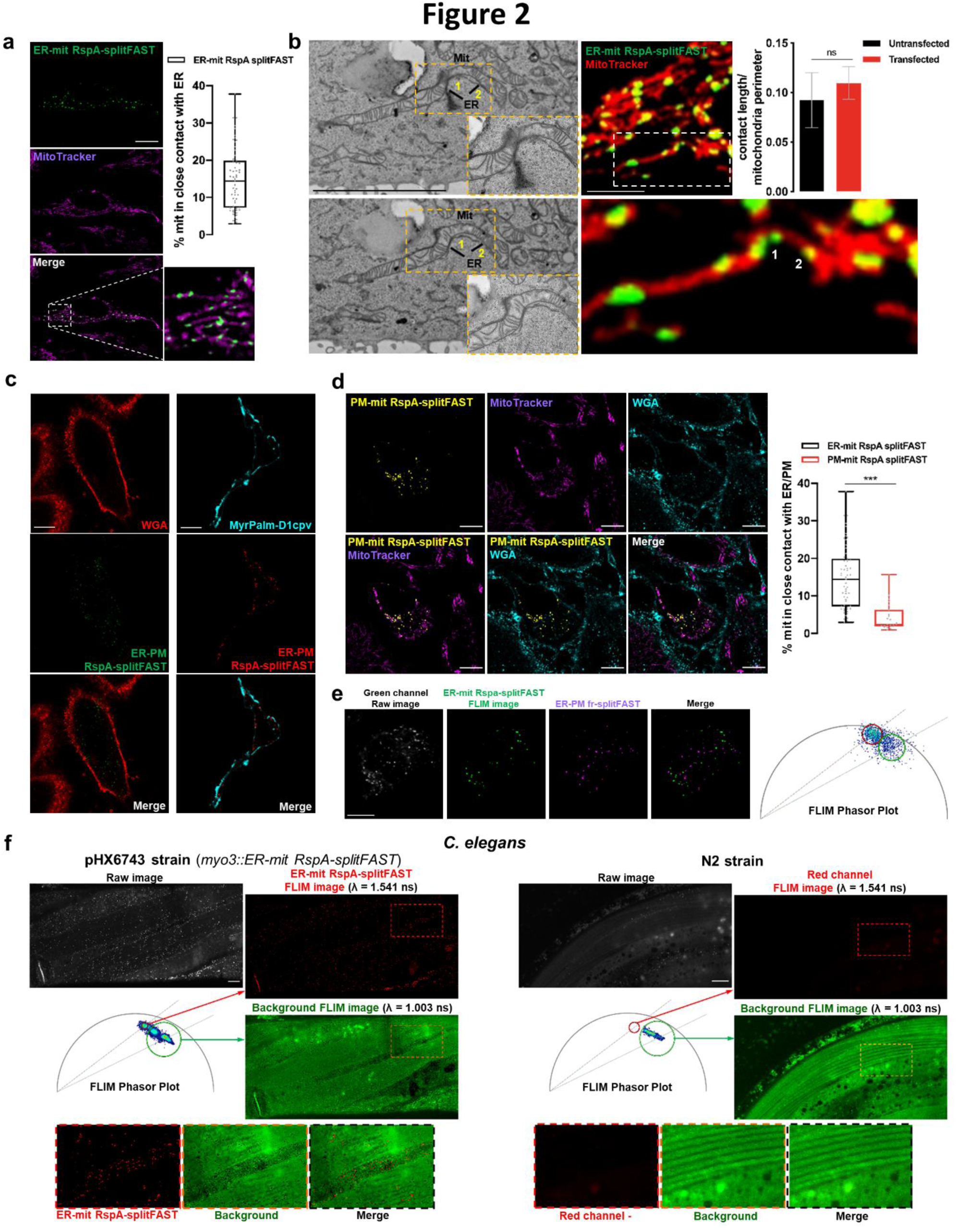
The RspA-splitFAST system to investigate ER-mit, ER-PM, PM-mit MCSs *in vitro* and *in vivo*. **a**, Representative confocal images of HeLa cells expressing the short ER-mit RspA-splitFAST and in which mitochondrial network was marked with MitoTracker Deep Red. On the right, the box plot shows the percentage of mitochondrial surface co-localized with ER-mit RspA-splitFAST. n=53 cells from 3 independent experiments. Scale bar, 10 µm. **b**, Representative EM and confocal images of the CLEM analysis showing a HeLa cell expressing the short ER-mit RspA-splitFAST probe and in which mitochondrial network was marked with MitoTracker Red. Fluorescent splitFAST dots (as an example, two dots were marked “1” and “2” in the figure) correspond to sites of close ER-mit membrane juxtaposition, as revealed by the two corresponding EM slices shown on the left (see also Supplementary Video 2). The expression of the probe does not force organelle tethering, as the fraction of mitochondrial perimeter in close contact with ER, analysed by CLEM, is similar in control (untransfected) and ER-mit RspA-splitFAST-expressing (transfected) cells. n=5 cells from 3 independent experiments. Scale bar, 5 µm. **c**, Representative confocal images of HeLa cells expressing ER-PM RspA-splitFAST. The probe marks ER-PM MCSs, while PM was labeled by either the chemical dye WGA or MyrPalm-D1cpv expression. ER-PM MCSs are visualized either in green or in red by addition of respectively HMBR or HBR-3,5DOM. Scale bar, 10 μm. **d**, Representative confocal images of PM-mit RspA-splitFAST-expressing HeLa cells. The probe marks mitochondrial-PM MCSs (yellow). Mitochondrial network (far-red) and PM (cyan) are labelled by MitoTracker and WGA, respectively. Scale bar, 10 μm. On the right, the box plots represent the percentage of mitochondrial surface engaged in the formation of close contacts with either ER (same box plot of panel **a**) or PM. n>23 cells from 3 independent experiments. p<0.001. **e**, Representative confocal images of HeLa cells co-expressing ER-PM fr-splitFAST and ER-mit RspA-splitFAST. While the far-red signal is exclusively emitted by ER-PM fr-splitFAST (unambiguously marking ER-PM MCSs), the green signal can be emitted by both ER-mit RspA-splitFAST and, to a lower level, ER-PM fr-splitFAST (note that fr-splitFAST can also incorporate the green and red fluorogens). As the lifetimes of the green signals emitted by respectively ER-mit RspA-splitFAST and ER-PM fr-splitFAST are different (see FLIM Phasor Plot on the right and Methods), we could distinguish ER-mit and ER-PM MCSs in the very same cells by FLIM. Scale bar, 10 μm. **f**, Representative confocal images of a *Caenorhabditis elegans* strain (pHX6743), expressing the ER-mit RspA-splitFAST under the *myo3* promoter (see Methods), to study ER-mit MCSs *in vivo*. The fluorescent signals of ER-mit RspA-splitFAST (red dots) and of background (BCKG) were separated by FLIM, exploiting their different lifetimes (See Phasor Plot; λ ∼1 ns for BCKG and ∼1.5 ns for ER-mit RspA-splitFAST). MCSs in body wall muscle cells (BWM) are shown. The control worm strain N2 (right) does not show any dotted staining in BWM cells after the incubation with the fluorogen.

Co-expression of ER-RspA-NFAST and PM-targeted RspA-CFAST (ER-PM RspA-splitFAST; **Supplementary Fig. 1e**) produced a dotted pattern corresponding to ER-PM MCSs (**Fig. 2c**), suggesting that this system can be tailored to detect different types of MCSs. By expressing this probe, in few cells we additionally observed some dots corresponding to ER-intracellular vesicles (likely early endosomes - EE) MCSs (**Fig. 2c**). This result, possibly associated with a partial endocytosis of PM-RspA-CFAST when highly expressed, supports the existence of close ER-EE MCSs, as previously reported^15,16^.

In addition, we co-expressed OMM-RspA-NFAST with PM-RspA-CFAST (PM-mit RspA-splitFAST) to detect PM-mit MCSs (**Fig. 2d**) and observed that a lower fraction of mitochondrial surface is in close contact with the PM (**Fig. 2d**), compared to that in contact with the ER (**Fig. 2a**). Moreover, by exploiting the spectral flexibility and versatility of splitFAST variants, the combined expression of PM-frNFAST, ER-RspA-CFAST and OMM-RspA-NFAST allowed us to simultaneously visualize ER-mit and ER-PM MCSs in the very same cells. Indeed, using fluorescence-lifetime imaging microscopy (FLIM), we could distinguish them by both their different spectral properties and fluorescence lifetime (**Fig. 2e**, see legend and Methods).

Finally, we generated a *C. elegans* strain expressing ER-mit RspA-splitFAST under the *myo3* promoter, to specifically visualize sarcoplasmic reticulum (SR)-mitochondria MCSs in body wall muscle cells (see Methods). By a FLIM-based approach, we could clearly identify SR-mitochondria MCSs in living worms separating them from background signal (**Fig. 2f**), suggesting these probes are suitable for *in vivo* investigations.

### The dynamics of ER-mit MCSs associate with ER, mitochondria and cytoskeleton remodelling

We followed organelle and MCS dynamics in live HeLa clones and COS-7 cells, expressing ER-mit RspA-splitFAST and stained for ER and/or mitochondrial markers. Imaging was performed by Airyscan-based super-resolution microscopy or lattice light-sheet microscopy (LLSM), allowing fast 4D acquisitions while minimizing photo-damage and preserving good spatial resolution. *Imaris*-based surface analysis (see Methods) revealed that the volume of most ER-mit RspA-splitFAST dots is < 0.1µm^3^, with the larger ones assuming an ellipsoidal shape (**Fig. 3a**). We observed that ER-mit MCSs are frequently associated with regions of mitochondrial branching and confirmed that they can be maintained during organelle movements^17^ (**Fig. 3b-c and Supplementary Videos 3-4**). Moreover, in line with previous findings, we noticed that they mark sites of dynamin-related protein 1 (DRP1)-puncta formation^18^ and mitochondrial fission^19^ (**Fig. 3d, Supplementary Fig. 2a and Supplementary Video 5**), but additionally observed frequent cases in which transient mitochondrial fusion (‘kiss-and-run’ mechanism)^20^ occurs at ER-mit MCSs (**Fig. 3e and Supplementary Video 6**). Importantly, by *Imaris*-based tracking, we noticed that several MCSs undergo themselves frequent fission/fusion events (**Fig. 3f and Supplementary Video 7**), tuning their continuous remodelling.

**Figure 3:**
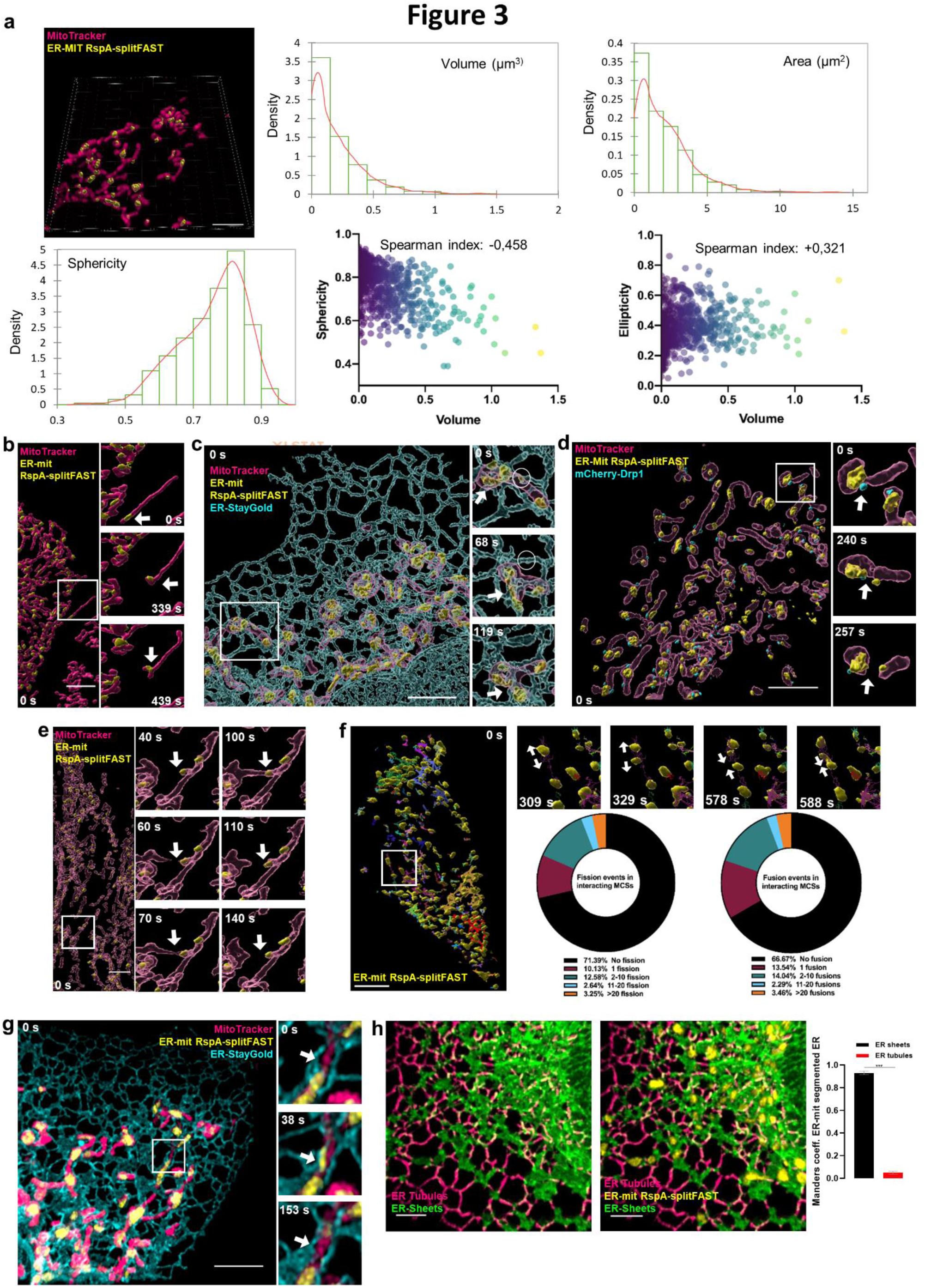
ER-mit MCS dynamics. **a,** Representative Airyscan 3D image of a COS-7 cell co-expressing the ER-mit RspA-splitFAST probe (surface rendering, yellow), the ER-StayGold (not shown) and stained with MitoTracker Deep Red (magenta). Scale bar, 5 μm. Distribution graphs represent the Kernel density (orange) and the histogram (green) of volume, area and sphericity values of ER-mit RspA-splitFAST dots, calculated using IMARIS (see Methods). n=10 cells from 3 independent experiments. Dispersion plots show the negative correlation (Spearman index: -0.458) between volume and sphericity (left) and the positive correlation (Spearman index: +0.321) between volume and ellipticity (right). **b,** Representative lattice light-sheet microscopy 3D image of a HeLa cell clone expressing the ER-mit RspA-splitFAST probe (surface rendering, yellow) and stained with MitoTracker Deep Red (surface rendering, magenta). Scale bar, 5 μm. Enlarged regions show the time-lapse of an extensive mitochondrial movement while an ER-mit MCS is maintained (arrow). In the last time frame (439 s), a mitochondrial branching at the contact site is visible (arrow). See also the corresponding Supplementary Video 3. **c,** Representative Airyscan image of a COS-7 cell as in panel (**a**) but with the ER-StayGold surface rendering also displayed (cyan). Scale bar, 5 μm. Enlarged regions show the time-lapse of a co-sliding event of mitochondria, ER and ER-mit RspA-splitFAST dots (arrow), as well as the disappearance of an ER-mit MCS (circle, first frame) upon mitochondria detachment from ER tubules. See also the corresponding Supplementary Video 4. **d,** Representative Airyscan 3D image of a COS-7 cell co-expressing ER-mit RspA-splitFAST (surface rendering, yellow), mCherry-Drp1 (surface rendering, cyan) and stained with MitoTracker Deep Red (surface rendering, magenta). mCherry-Drp1 signal localizes in proximity of ER-mit MCSs. Scale bar, 5 μm. Enlarged regions show the time-lapse of mCherry-Drp1 localization at the site of mitochondria (as well as of ER-mit MCS) fission (arrow). See also the corresponding Supplementary Video 5. **e,** Representative image as for panel (**b**). Scale bar, 5 μm. Enlarged regions show the time-lapse of an ER-mit MCS movement (arrow) at a site of mitochondria “Kiss-and-run” fusion (40-70 s) followed by organelle fission (100-140 s). See also the corresponding Supplementary Video 6. **f,** Representative 3D image of a HeLa cell expressing the ER-mit RspA-splitFAST probe (surface rendering, yellow). Scale bar, 5 μm. ER-mit RspA-splitFAST dots were tracked using IMARIS (see Methods) along the mitochondrial network stained with MitoTracker Deep Red (not shown). ER-mit MCSs move in space and interact each other. Tracks (depicted as continuous lines) report MCS movements and interactions. Each track is represented in a different color. If two ER-mit MCSs undergo fusion or fission, they are considered part of the same track. Enlarged regions show the time-lapse of a MCS undergoing fission (309-329 s), and two MCSs fusing together (578-588 s). Charts represent the percentage of interacting ER-mit MCS subgroups (tracks) for each range of fission/fusion events, as indicated. n= 4 cells from 2 independent experiments. See also the corresponding Supplementary Video 7. **g,** Representative Airyscan 3D image of a COS-7 cell co-expressing the ER-mit RspA-splitFAST probe (yellow), ER-StayGold (cyan) and stained with MitoTracker Deep Red (magenta). Scale bar, 5 μm. Enlarged regions show the time-lapse of the appearance (38 s) of an ER-mit MCS (arrow), upon ER and mitochondria membrane docking, and its following disappearance (153 s), upon organelle membrane distancing (arrow). See also the corresponding Supplementary Video 8. **h,** Representative 3D image of a COS-7 cell co-expressing the ER-mit RspA-splitFAST probe (yellow), ER-StayGold and stained with MitoTracker Deep Red (not shown). ER-StayGold signal has been segmented (see Methods) to separate ER tubules (magenta) from ER sheets (green). On the right, Manders’ co-localization M2 coefficient quantification shows that the majority of short ER-mit MCSs colocalize with ER sheets. Mean ± SEM; n=5 cells from 3 independent experiments. Scale bar = 5 µm.

Moreover, Airyscan-based analysis revealed that the transient docking of ER tubules to mitochondria is promptly matched with the appearance and vanishing of ER-mit RspA-splitFAST fluorescent dots (**Fig. 3g and Supplementary Video 8**), confirming that the probe is dynamic and fully reversible. Interestingly, in COS-7 cells, we found that the majority of short ER-mit MCSs involves ER sheets, rather than tubules (**Fig. 3h**), though the remodelling of ER tubules frequently underlies ER-mit MCS fusion/fission (Supplementary Fig. 2b **and Supplementary Video 9**). Finally, we observed that most ER-mit MCSs associate with tubulin- and F-actin-positive structures (**Supplementary Fig. 2c and Supplementary Video 10**), in line with the key role of different cytoskeleton elements in organelle dynamics and MCS rearrangement^17,18,20^.

### Detection of MCS perturbations in Alzheimer’s disease (AD) cell models by splitFAST-based probes

We and others have previously demonstrated that ER-mitochondria association is strengthened in different AD models^3,21–23^. To test whether ER-mit RspA-splitFAST can efficiently detect these alterations, we firstly confirmed it can be used in AD-relevant primary cells, including microglia (**Fig. 4a**), astrocytes (**Fig. 4b**) and cortical neurons (**Fig. 4c-d**), in which we observed that ER-mit MCSs distribute throughout the soma, axons and dendrites (**Fig. 4d and Supplementary Video 11**).

**Figure 4:**
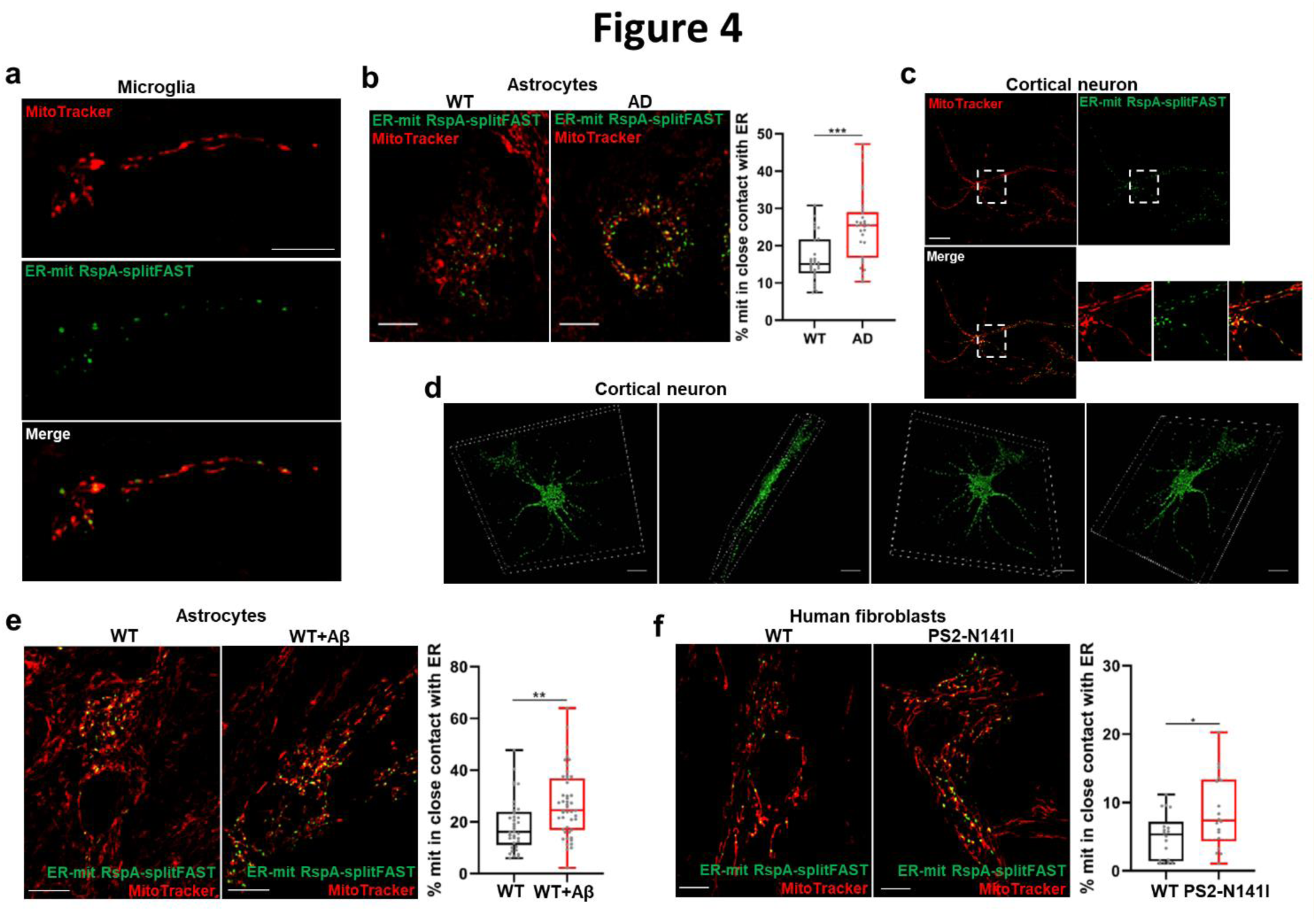
ER-mit MCSs in different brain cells and Alzheimer’s disease cell models. **a-f**, Representative confocal images of different ER-mit RspA-splitFAST-expressing cells. Where indicated, cells were co-stained with MitoTracker Deep Red. Scale bar, 10 µm. ER-mit MCSs are shown in mouse primary microglia (**a**), astrocytes (**b, e**), cortical neurons (**c, d**) and human fibroblasts (**f**). In **b**, **e, f**, the box plots represent the percentage of mitochondrial surface co-localized with ER-mit RspA-splitFAST for the indicated cell types. In **b**, astrocytes from WT or AD mice were compared. n>30 cells from 3 independent experiments. In **e**, astrocytes from WT mice were exposed (WT + Aβ) or not (WT) to a conditioned medium containing naturally generated Aβ peptides (see Methods). n>37 cells from 3 independent experiments. In **f**, primary human skin fibroblasts from either a healthy donor (WT) or a familial Alzheimer’s disease patient (PS2-N141I) were compared. n>18 cells from 3 independent experiments. *p<0.05; **p<0.01; ***p<0.001. **d**, 3D projection of the ER-mit RspA-splitFAST fluorescent signal in a cortical neuron, imaged by confocal microscopy. Scale bar, 10 µm. See also the corresponding Supplementary Video 11.

As expected, astrocytes from an AD transgenic mouse model (B6.152H)^24^ displayed a significantly higher fraction of mitochondrial surface covered by the ER-mit RspA-splitFAST signal, compared to wild type (WT) cells (**Fig. 4b**). A similar increase in ER-mit MCSs was detected in astrocytes from WT mice upon acute exposure to naturally generated amyloid β (Aβ) peptides (**Fig. 4e**; see Methods), suggesting that multiple pathways involved in AD might converge on alterations of ER-mitochondria juxtaposition. Similarly, a higher ER-mitochondria tethering was observed in human fibroblasts from a patient harboring a familial AD mutation in Presenilin 2, compared to those from an age-matched healthy donor (**Fig. 4f**), as previously reported^23,25,26^. Overall, these data demonstrate that the splitFAST-based probes can detect MCS alterations in disease models.

### PRINCESS design and characterization

Several cell pathways are regulated by Ca^2+^ signalling events occurring at MCSs. For instance, during inositol 1,4,5-trisphosphate (IP3)-linked cell stimulations, transient microdomains of high Ca^2+^ concentration are formed at the ER-mitochondria interface, sustaining mitochondrial Ca^2+^ uptake^27,28^. Similarly, upon depletion of ER Ca^2+^ content, the activation of store-operated Ca^2+^ entry (SOCE) at ER-PM MCSs is instrumental in recovering ER Ca^2+^ content^29^.

However, the detection of these rapid and spatially restricted Ca^2+^ hot spots has been challenging. Indeed, the limitations of current techniques, mostly relying on slow super-resolution microscopy or plagued by limited spatial resolution and complex pixel-by-pixel analyses^30,31^, hinder the possibility of unambiguously locating the signal of interest to a specific MCS.

We tackled this problem using two different approaches. First, we co-expressed ER-mit RspA-splitFAST (to identify ER-mit MCSs) with a reported Ca^2+^ probe (Cepia3)^32^, that we targeted to the whole OMM (**Fig. 5a**). Importantly, Cepia3 was chosen for this experiment because its Ca^2+^ affinity (K_d_ ∼6-7 µM; **Fig. 5b**) well-suits the reported cation concentration reached at ER-mitochondria interface (∼10-15 µM)^30,31^. IP3-linked cell stimulations induced an increase of OMM-Cepia3 ratio (**Fig. 5a**; see Methods), associated with ER Ca^2+^ release and cytosolic Ca^2+^ rise experienced by the OMM. However, by restricting the analysis to the sub-regions of OMM-Cepia3 co-localized with ER-mit RspA-splitFAST signal, larger and faster increases in Cepia3 ratio were observed (**Fig. 5a**), confirming that ER-mit MCSs are sites of high Ca^2+^-microdomain generation and privileged inter-organelle Ca^2+^ transfer.

**Figure 5:**
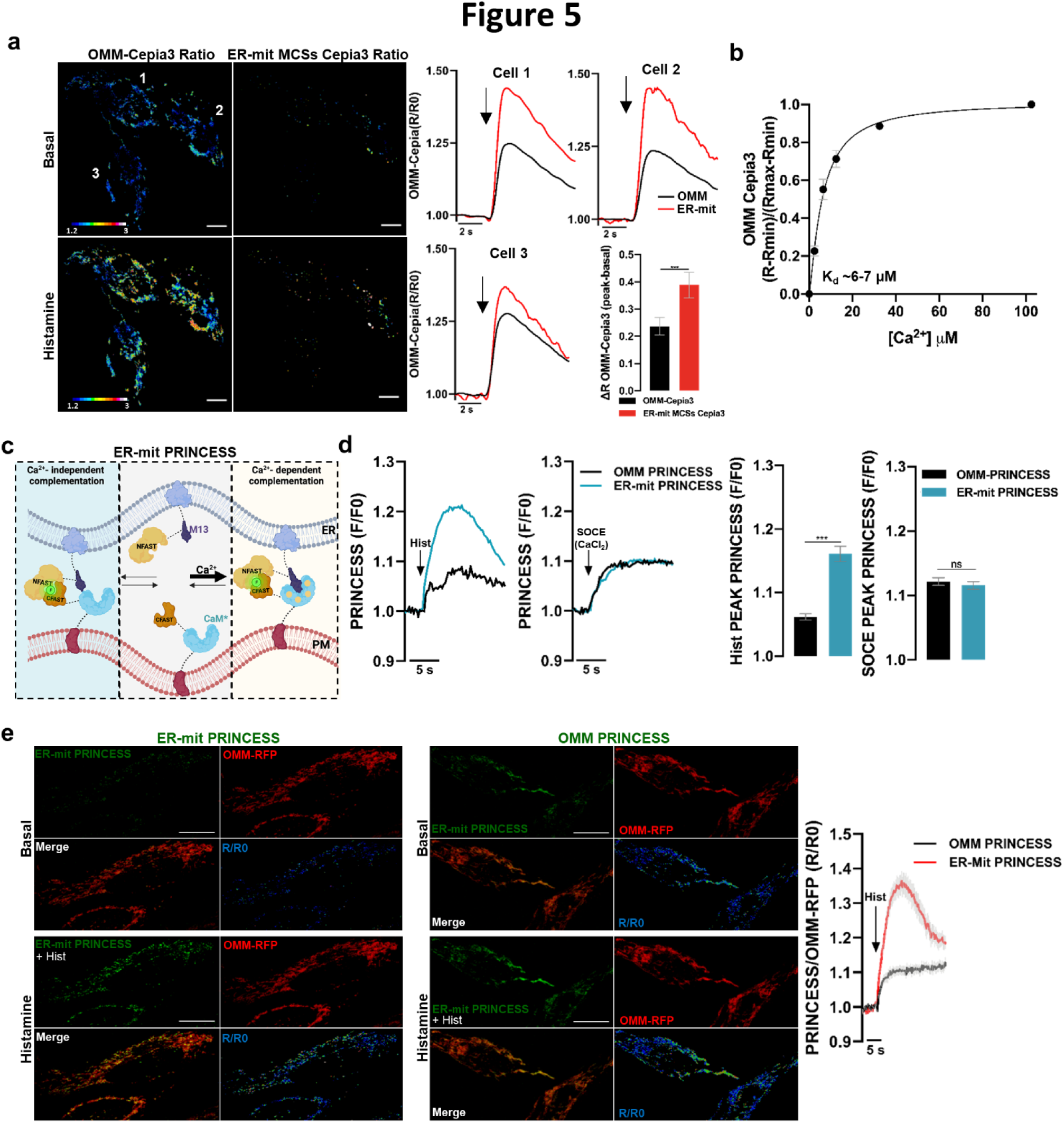
Measurements of Ca^2+^ signals at MCSs by PRINCESS. **a,** Representative images of the 475/390 nm ratio (R, see Methods) of OMM-Cepia3, co-expressed in HeLa cells with ER-mit RspA-splitFAST (not shown) to mark ER-mit MCSs. The ratio images of OMM-Cepia3 and of the portion of OMM-Cepia3 co-localized with ER-mit RspA-splitFAST (ER-mit MCSs Cepia3 Ratio) are shown, before (basal) and upon histamine (100 µM) stimulation. On the right, traces represent the ratios of OMM-Cepia3 (OMM) or of the portion of OMM-Cepia3 co-localized with ER-mit RspA-splitFAST (ER-mit) upon histamine stimulation (arrows), for the three cells on the left. Bars represent the ΔR upon histamine stimulation for the indicated conditions. Mean ± SEM, n=9 cells from 3 independent experiments. ***p<0.001. **b,** OMM-Cepia3, expressed in HeLa cells, was calibrated by permeabilizing cells with digitonin (25 µM) in intracellular-like buffer and adding fixed Ca^2+^ concentrations. Ratiometric 475/390 nm measurements were performed (see Methods). Mean ± SEM, n=12 cells from 4 independent experiments. **c,** The cartoon represents the rationale behind ER-mit PRINCESS design. Mutated Calmodulin (CaM*) from Cepia3 and the M13 peptide were incorporated within ER-mit splitFAST, to provide Ca^2+^-sensing capabilities. **d,** Representative traces of ER-mit PRINCESS or OMM-PRINCESS fluorescence increases in HeLa cells, upon histamine (100 µM) stimulation in Ca^2+^-free mKRB (see Methods), or CaCl_2_ (2 mM) addition (SOCE) after 6 min depletion of ER Ca^2+^ content (obtained by histamine and thapsigargin (100 nM) stimulation in Ca^2+^-free mKRB). On the right, bars represent the peaks of OMM- or ER-mit PRINCESS fluorescence (expressed as F/F0) upon the indicated treatments. Mean ± SEM; n>27 cells from of at least 3 independent experiments. ***p<0.001. **e,** Representative confocal images acquired at high speed with resonant scanner (see Methods) of HeLa cells co-expressing OMM-RFP with either ER-mit PRINCESS or OMM-PRINCESS, before (basal) or upon histamine (100 µM) stimulation in Ca^2+^-free mKRB, as indicated. The images of the ratio between PRINCESS and OMM-RFP fluorescent signals are also shown. These images are frames of the corresponding Supplementary Videos 12 and 13. On the right, the mean traces ± SEM of the ratio between each specific PRINCESS and OMM-RFP fluorescent signals upon histamine stimulation are shown, as indicated. n=19 (OMM PRINCESS) and 27 (ER-mit PRINCESS) cells from 3 independent experiments.

In these experiments, the co-expression of ER-mit RspA-splitFAST is helpful to increase the specificity of the analysis, allowing to simultaneously visualize ER-mit MCSs in a different colour. However, though it represents an improvement, this approach does not completely overcome the limitations of previous techniques, because it requires the expression of two different sensors (*i.e.*, ER-mit RspA-splitFAST and the OMM-Cepia3 Ca^2+^ probe) and the co-localization of their different fluorescent signals, inevitably retaining some resolution constraints.

An ideal method would be that of unambiguously marking MCSs and specifically measuring the associated Ca^2+^ dynamics by a single sensor, endowed with sufficient spatiotemporal resolution to report local, fast changes of Ca^2+^ concentration. To design such a probe, that we dubbed PRINCESS, we integrated into a single reporter both splitFAST (tailored to detect MCSs) and a couple of known Ca^2+^-sensing protein-domains (*i.e.*, Calmodulin -CaM- and the M13 peptide), thus endowing it with Ca^2+^-detection capabilities (**Fig. 5c**). Importantly, to match the range of Ca^2+^ concentrations potentially reached at MCSs, we selected the low Ca^2+^ affinity CaM mutant (CaM*) of Cepia3^32^ (**Fig. 5b**), which should restrict the increase in splitFAST complementation only where Ca^2+^ hotspots are generated. We hypothesized that, in resting conditions, PRINCESS should maintain a low-level, spontaneous NFAST-CFAST complementation at MCSs, allowing to mark organelle contacts. However, upon sustained Ca^2+^ peaks, the transient CaM*-M13 binding could dynamically change the equilibrium of the reaction, favouring an interaction-dependent complementation of a larger fraction of NFAST-CFAST complexes (**Fig. 5c**).

We generated different versions of this probe, to specifically investigate Ca^2+^ dynamics at ER-mit (ER-mit PRINCESS; **Fig. 5c**) or ER-PM (ER-PM PRINCESS; **Supplementary Fig. 3a**) MCSs. HeLa cells expressing ER-mit PRINCESS displayed a dotted pattern along the mitochondrial network (**Supplementary Fig. 3b**), like that observed with the ER-mit RspA-splitFAST (**Fig. 2a**) and corresponding to ER-mit MCSs. Notably, IP3-linked cell stimulations triggered a fast, transient increase of ER-mit PRINCESS fluorescence (**Fig. 5d-e**). Conversely, the activation of Ca^2+^ entry through the PM by SOCE, which does not generate microdomains of high Ca^2+^ concentration at ER-mit MCS^30^, led to a slower and lower fluorescence rise (**Fig. 5d**). These data suggest that the probe can promptly detect Ca^2+^ peaks generated at ER-mitochondria interface upon specific cell stimulations. To further verify this point, we compared the response of ER-mit PRINCESS with that of a similar, control probe, obtained by targeting both the M13-NFAST and CaM*-CFAST fragments to the whole OMM (OMM PRINCESS; **Fig. 5d**). The prediction is that OMM PRINCESS should detect Ca^2+^ rises occurring along the entire OMM, and not specifically at ER-mit MCSs. Upon histamine-induced, IP3-dependent Ca^2+^ rises, the response of ER-mit PRINCESS was faster and higher than that of OMM PRINCESS, whereas the two probes displayed a similar behaviour upon SOCE activation (**Fig. 5d**). Importantly, by co-expressing ER-mit or OMM PRINCESS together with an OMM-RFP, we performed ratiometric measurements that confirmed a higher response of ER-mit PRINCESS upon histamine-induced cell stimulations (**Fig. 5e and Supplementary Videos 12-13**).

Interestingly, when similar experiments were performed with ER-PM PRINCESS, we observed comparably high and fast fluorescence increases upon Ca^2+^ rises triggered either by histamine or SOCE (**Supplementary Fig. 3c**). Therefore, ER-PM MCSs experience high Ca^2+^ concentrations upon both types of cell stimulation. This is in line with the reported intense IP3-receptor activity^33^, as well as the known activation of Orai channels during SOCE^29^, at ER-PM junctions.

### ER-PM and ER-mitochondria MCSs are remodelled by Ca^2+^ signals

The data above confirm previous findings indicating that MCSs host specific Ca^2+^ signalling events. By taking advantage of the dynamic complementation of splitFAST-based probes, we wondered whether the opposite is also true; *i.e.*, whether Ca^2+^ signals in turn modulate organelle juxtaposition. Total internal reflection fluorescence (TIRF)-based experiments have corroborated a Ca^2+^-mediated regulation of ER-PM MCSs^34^; however, studying the possible existence of similar mechanisms for MCSs involving organelles located far away from the PM has been challenging.

First, we checked whether ER-PM RspA-splitFAST is sensitive enough to detect the reported strengthening of ER-PM junction upon release of ER Ca^2+^ content, associated with the recruitment of the ER transmembrane proteins Stromal Interaction Molecule 1/2 (STIM1/STIM2) at ER-PM MCSs and activation of SOCE^29,35^. We indeed observed a progressive increase of ER-PM RspA-splitFAST signal, reaching a plateau after ∼10 minutes (**Supplementary Fig. 4a**). Importantly, by tailoring RspA-splitFAST to specifically investigate the migration of STIM1 close to PM, we found that this process precedes the rise of ER-PM juxtaposition and is transient (**Supplementary Fig. 4a**), confirming that STIM1 recruitment is critical for triggering the formation/expansion of ER-PM MCSs, but that additional proteins contribute to their stabilization.

As to ER-mit MCSs, their architecture is critical for the formation of Ca^2+^ microdomains facilitating inter-organelle Ca^2+^ transfer, as confirmed by ER-mit PRINCESS. However, whether and how these signals might, in turn, modulate ER-mitochondria juxtaposition has been poorly investigated. In HeLa cells, we did not observe rapid changes of the ER-mit RspA splitFAST signal upon histamine-elicited elevations of cytosolic Ca^2+^ (**Supplementary Fig. 4b and Supplementary Video 14**), implying that transient Ca^2+^ peaks do not directly affect ER-mit MCSs. However, we noted a tendency of the fluorescent signal to slightly and progressively increase after few minutes following drug addition (**Fig. 6a**, black trace), an effect that was enhanced by sustaining histamine-induced Ca^2+^ peaks via cell co-stimulation with thapsigargin (Tg), a SERCA pump inhibitor (**Fig. 6a**, red trace). To verify whether long-lasting cytosolic Ca^2+^ elevations could underlie ER-mit MCS potentiation, we pre-incubated cells with BAPTA-AM, a known intracellular Ca^2+^ buffer. Surprisingly, we observed an even higher increase in ER-mit RspA splitFAST signal (*i.e.*, in ER-mitochondria juxtaposition) upon histamine-Tg exposure (**Fig. 6a**, green trace), suggesting that cytosolic Ca^2+^ is not involved. Moreover, in this condition, we observed that the signal of the probe was slightly increasing ahead of cell stimulation (**Fig. 6a**, green trace). We reasoned that this effect could be linked to a partial depletion of ER Ca^2+^ content during cell pre-incubation with BAPTA-AM, as previously reported^36^ and here confirmed (**Supplementary Fig. 4c**). Moreover, the presence of BAPTA-AM remarkably accelerates the discharging of ER Ca^2+^ upon histamine-Tg stimulation (**Supplementary Fig. 4c)**, possibly explaining the observed faster increase in ER-mitochondria juxtaposition (**Fig. 6a**). To assess whether ER Ca^2+^ depletion potentiates ER-mit MCSs, we directly and acutely buffered Ca^2+^ in the lumen of the ER by TPEN, a low-affinity Ca^2+^ chelator that specifically sequesters Ca^2+^ within the ER (**Supplementary Fig. 4d**) whilst not interfering with cytosolic Ca^2+^ rises^37^. Acute TPEN addition progressively increases the ER-mit RspA-splitFAST signal (**Fig. 6a**, blue trace); along the same line, different treatments partially reducing ER Ca^2+^ content (*i.e.*, downregulation of SERCA pump, TPEN treatment or incubation with BAPTA-AM; **Supplementary Fig. 4c-e**) enhance the fraction of mitochondrial surface engaged in the formation of MCSs with the ER (**Supplementary Fig. 4f**). Remarkably, we observed that this effect is reversible because the refilling of ER Ca^2+^ content restored basal levels of ER-mit MCSs, though the recovery of organelle tethering was delayed of several minutes (**Fig. 6b**). Overall, these data support a hitherto unknown mechanism, in which the content of Ca^2+^ in the lumen of the ER modulates ER-mitochondria coupling.

**Figure 6:**
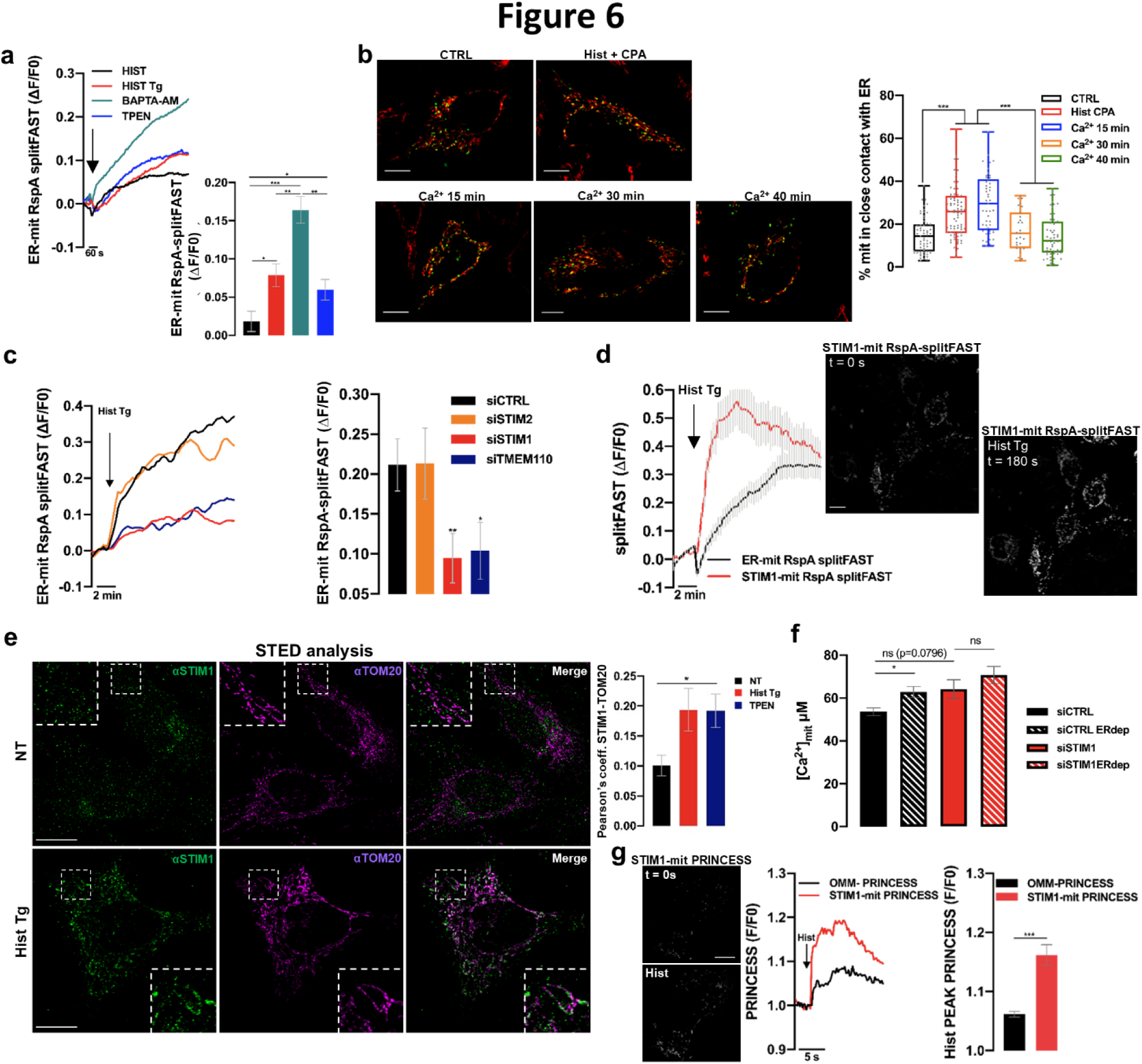
STIM1 mediates ER-mit MCS remodelling upon ER Ca^2+^ depletion. **a**, Representative traces of ER-mit RspA-splitFAST fluorescence in HeLa cells upon ER Ca^2+^ depletion by different treatments (arrow: 100 μM histamine, HIST; 100 μM histamine plus 100 nM thapsigargin (HIST Tg); pre-treatment with 10 µM BAPTA-AM for 30 minutes followed by 100 μM histamine plus 100 nM thapsigargin (BAPTA-AM); 500 μM TPEN). On the right, bars represent the increase in ER-mit RspA-splitFAST fluorescence (expressed as ΔF/F0) 15 min after the indicated cell stimulations. Mean ± SEM; n>49 cells from 4 independent experiments. *p<0.05; **p<0.01; ***p<0.001. **b**, Representative confocal images of HeLa cells, expressing the ER-mit RspA-splitFAST probe and labelled with MitoTracker, in different conditions (at basal, CTRL; 15 min after stimulation with 100 μM histamine plus 20 μM cyclopiazonic acid, Hist CPA; and 15, 30, or 40 min after the removal of the Hist + CPA stimulus and re-addition of 2 mM CaCl_2_, to allow ER Ca^2+^ refilling). Scale bar, 10 μm. On the right, the box plots represent the percentage of mitochondrial surface (% mit) engaged in close contact with the ER for each condition. n>27 cells from 3 independent experiments. **c**, Representative traces of ER-mit RspA-splitFAST fluorescence upon ER Ca^2+^ depletion by 100 μM histamine plus 100 nM thapsigargin (Hist Tg) in HeLa cells transfected with control siRNA (siCTRL, black trace) or knocked down by specific siRNAs for STIM1 (siSTIM1, red trace), STIM2 (siSTIM2, orange trace) or TMEM110 (siTMEM110, blue trace). On the right, bars represent the increase in ER-mit RspA-splitFAST fluorescence (expressed as ΔF/F0) after 15 min from cell stimulation, for the indicated conditions. Mean ± SEM; n>22 cells from 3 independent experiments. *p<0.05; **p<0.01. **d**, Mean ± SEM traces of either ER-mit RspA-spiltFAST or STIM1-mit RspA-splitFAST fluorescent signal upon ER Ca^2+^ depletion (by 100 μM histamine plus 100 nM thapsigargin; Hist Tg) in HeLa cells, as indicated. n>56 cells from 3 independent experiments. On the right, representative images of HeLa cells expressing STIM1-mit RspA-splitFAST, before (t = 0 s) or 180 s after exposure to Hist plus Tg, as indicated. **e**, Representative STED images of HeLa cells unstimulated (NT) or stimulated for 15 min with 100 μM histamine plus 100 nM thapsigargin (Hist Tg, to induce ER Ca^2+^ depletion), and immunolabeled with αSTIM1 (green) and αTOM20 (magenta) antibodies. Magnifications show regions where the recruitment of STIM1 close to mitochondria upon ER Ca^2+^ depletion is more evident. Scale bar, 10 μm. On the right, bars represent the Pearson’s co-localization coefficient between endogenous STIM1 and TOM20 signals, after 15 min in these two conditions (NT and Hist Tg) or upon 500 μM TPEN treatment (as an alternative treatment to induce ER Ca^2+^ depletion). Mean ± SEM; n>40 cells from 3 independent experiments. *p<0.05. **f**, Bars represent mitochondrial Ca^2+^ peaks observed upon IP3-induced ER Ca^2+^ release in Hela cells, expressing the Ca^2+^ sensor Aequorin in the mitochondrial matrix (see Methods) and transfected with specific siRNA for STIM1 (siSTIM1) or control siRNA (siCTRL). Before the experiments, cells were treated (ERdep) or not with 500 μM TPEN (15 min) to induce partial ER Ca^2+^ depletion and thus the remodelling of ER-mit MCSs, followed by removal of TPEN just before the aequorin experiments, to allow a fast re-filling of ER Ca^2+^ content. Mean ± SEM; n>11 independent experiments. *p<0.05. **g**, Representative traces of PRINCESS fluorescence in OMM-PRINCESS- or STIM1-mit PRINCESS-expressing HeLa cells, treated with 100 μM histamine (Hist) in Ca^2+^-free, EGTA containing mKRB. On the right, bars represent the peaks of OMM- or STIM1-mit PRINCESS fluorescence (expressed as F/F0) upon histamine stimulation. Data of OMM-PRINCESS are the same of Fig. 5d, here reported for a better comparison. Mean ± SEM; n>28 cells from 3 independent experiments. ***p<0.001. On the left, representative fluorescent images of HeLa cells expressing STIM1-mit PRINCESS, before (t = 0 s) and 2 s after stimulation with histamine (Hist).

### STIM1 is involved in the Ca^2+^-mediated modulation of ER-mit MCSs

Using ER-mit RspA-splitFAST, we explored the molecular mechanism of the Ca^2+^-mediated modulation of ER-mit MCSs by a siRNA-based screening of some possible protein effectors, chosen because previously reported to be Ca^2+^-sensitive and/or involved in the modulation of different MCSs (**Supplementary Table 1**). Among these proteins, we excluded an involvement of PERK (<u>P</u>rotein Kinase RNA-Like <u>ER</u> <u>K</u>inase), previously shown to strengthen ER-PM juxtaposition by a Ca^2+^-dependent mechanism^38^. Indeed, though PERK downregulation reduces ER-mit MCSs in resting conditions (**Supplementary Fig. 5a**), it did not significantly affect their increase following the depletion of ER Ca^2+^ content (**Supplementary Fig. 5b**), not was this increase altered in PERK knockout (PERK-KO) cells (**Supplementary Fig. 5c**).

Conversely, we observed that the downregulation of STIM1, but not of STIM2, dampens the increase of ER-mitochondria juxtapositions upon release of ER Ca^2+^ (**Fig. 6c**). Intriguingly, a similar effect was detected by downregulating the STIM1 partner TMEM110 (Transmembrane protein 110; **Fig. 6c**)^39^, strongly supporting a possible involvement of this pathway in ER-mit MCS remodelling.

Experiments performed in SH-SY5Y cells double KO for both STIM1 and STIM2 (STIM1/2-dKO)^40^ confirmed a significant reduction of the effect of ER Ca^2+^-depletion on ER-mit MCSs, promptly rescued by the re-expression of STIM1 alone (**Supplementary Fig. 5d**). Next, we tailored RspA-splitFAST to test whether STIM1, upon sensing the depletion of ER Ca^2+^ content by its luminal EF-hand domains, migrates at ER-mit MCSs (**Supplementary Fig. 5e**), as it does towards ER-PM MCSs (**Supplementary Fig. 4a**)^29^. We indeed observed such a recruitment, preceding the rise of ER-mitochondria juxtaposition (**Fig. 6d**). Importantly, using super-resolution STED microscopy, we confirmed that a fraction of endogenous STIM1 is recruited and forms puncta close to mitochondria following ER Ca^2+^ depletion (**Fig. 6e**), excluding possible artifacts linked to its overexpression. Next, we wondered whether STIM1 can also act as a regulator of ER-mitochondria tethering in resting conditions (*i.e.*, without reducing ER Ca^2+^ content). Our results showed that the fraction of mitochondrial surface in contact with ER is not affected by STIM1 downregulation (**Supplementary Fig. 5f**), implying a specific role of this protein in boosting ER-mit MCSs only upon ER Ca^2+^-depletion.

We speculated that, upon reduction of ER Ca^2+^ concentration, the STIM1-mediated strengthening of organelle connectivity might be instrumental for sustaining physiological, constitutive levels of ER-to-mitochondria Ca^2+^ transfer, in turn critical to stimulating mitochondrial activity and cell bioenergetics^41,42^. In line with this hypothesis, after a transient TPEN-mediated partial depletion of ER Ca^2+^ content (see Methods), we measured a higher efficiency of ER-mitochondria Ca^2+^ shuttling in controls, but not in cells in which STIM1 was downregulated (**Fig. 6f**). This suggests that STIM1 might be specifically involved in the remodelling of those ER-mit MCSs hosting Ca^2+^ transfer activity, *i.e.*, where Ca^2+^ hot spots are generated. Indeed, by designing a PRINCESS probe to monitor Ca^2+^ dynamics where STIM1 is in contact with mitochondria (STIM1-mit PRINCESS; **Supplementary Fig. 5g**), we observed that these regions experience microdomains of high Ca^2+^ concentration upon IP3-linked cell stimulations (**Fig. 6g**). Overall, these data support a role of STIM1 in the modulation of ER-mitochondria Ca^2+^ shuttling at MCSs.

## Discussion

By tailoring specific variants of splitFAST endowed with low self-complementation, we here designed several chemogenetic reporters to dynamically investigate different MCSs and associated signalling. As in the case of previously reported bimolecular fluorescent complementation (BiFC) approaches, these reporters ensure high and tunable spatial resolution, thanks to an interaction-dependent generation of the fluorescent signal. However, compared with other BiFC systems, the incorporation of improved splitFAST versions guarantees a rapid and fully reversible complementation of the split fragments. This key property endows these reporters with unprecedented spatiotemporal resolution, enabling real-time monitoring of dynamic MCS changes, whilst minimizing the possible impact on physiological MCS rearrangements. Moreover, their wide spectral flexibility allows the combination with different probes for performing multi-colour imaging in the very same cell, while the small size of the FAST fragments reduces the risk of dysfunctional protein fusions.

The modest affinity of NFAST and CFAST fragments leads to a low-level, promptly reversible FAST self-complementation, which involves only a minor fraction of the molecules^9,14^. This implies that, depending on the sensitivity of the detection method, relatively high expression levels of the probe might be necessary to guarantee a sufficiently bright fluorescent signal at the MCSs under investigation. The fluorogens also contribute to increasing NFAST/CFAST affinity^14^; thus, the optimal fluorogen concentration should be tested in each experimental setup. We recommend using low fluorogen concentration (< 3-5 µM), avoiding excessive amounts to minimize the possible impact on MCS dynamics and reduce background. By generating a *C. elegans* strain and substantially improving the signal-to-noise ratio by FLIM, we demonstrated that these reporters can efficiently detect MCSs *in vivo*. We envisage that *in vivo* experimental setups will benefit from the availability of the far-red emitting fr-splitFAST (**Fig. 1h**), enabling a deeper tissue imaging and reducing phototoxicity.

By taking advantage of the dynamic and reversible complementation of our probes, we interrogated the relationship between Ca^2+^ signalling and MCSs, identifying a hitherto unknown pathway in which ER Ca^2+^ content modulates ER-mitochondria juxtaposition. Intriguingly, by an siRNA-based screening, we identified STIM1 as a key protein involved in this regulation. While the precise definition of the underlying molecular mechanism will require additional investigations, we further benefited from the versatility of these chemogenetic reporters, generating specific probes that allowed us to demonstrate the prompt recruitment of STIM1 at ER-mit (as well as ER-PM) MCSs upon ER Ca^2+^ discharge.

The intrinsic dynamicity of MCSs makes cell imaging the ideal application for our reporters; however, the built-in fluorogenic properties provided by splitFAST might enable to carry out larger screening by techniques such as cytometry. As a hint of this possible approach, we used fluorescence-activated cell sorting (FACS) to select the cell clones expressing ER-mit splitFAST (**Supplementary Fig. 1c**).

Last, by integrating Ca^2+^-sensing capabilities with both the high spatiotemporal resolution and the fluorescence activation provided by splitFAST, we introduced PRINCESS. In this innovative series of reporters, the spontaneous, low-level NFAST/CFAST self-complementation can be transiently boosted by the Ca^2+^-mediated binding of CaM* to M13, which favours a fully reversible interaction of the two fragments. We demonstrated that the unprecedented properties of PRINCESS allow to simultaneously detect MCSs and measure the associated Ca^2+^ dynamics using a single probe. We envisage that this approach will pave the way to develop an innovative class of biosensors, enabling multiple parameters to be monitored at MCSs and addressing a large diversity of unanswered biological questions.

## Methods

### Cell culture and transfection

HeLa (ATCC CCL-2) and COS-7 (ATCC CRL-1651) cells were maintained in DMEM (Sigma-Aldrich, D5671) supplemented with 10% FCS, 2 mM L-glutamine, 100 U/ml penicillin and 0.1 mg/ml streptomycin. Cells were growth in a humidified Heraeus incubator at 37°C, with 5% di CO_2_.

Tetracyclin-Regulated Expression (T-Rex) HeLa cells were maintained as previously described^43^. The expression of ER-mit splitFAST in the different clones was induced by addition of tetracyclin (2 µg/ml) for 16 h.

SH-SY5Y WT or STIM1/STIM2-dKO cells were previously described^40^ and maintained in DMEM (Gibco 41966-029) supplemented with 10% FCS and 1% MEM NEAA (Gibco 11140-035).

HeLa cells PERK-KD (shP) and relative controls (EV), as well as MEF WT and PERK-KO, were maintained as previously described^38^.

Transfection was performed at 50-60% confluence, 24-48 h after seeding, with either TransIT-LT1 (Mirus Bio, HeLa cells) or LipofectamineTM 2000 (Thermo Fisher, COS-7 and SH-SY5Y cells). Imaging was performed 24 h after transfection. Co-transfection of siRNA was performed with TransIT-X2 (Mirus Bio) and experiments perfomed 48 h later.

Transduction was performed with lentiviruses (Vector Builder, see plasmids) in medium containing polybrene (1 µg/ml). 24 h later, the medium was replaced with fresh medium. Imaging (or antibiotic selection) was performed at least 72-96 h after transduction.

### Primary cell cultures

Primary human skin fibroblasts from a FAD-PS2-N141I female patient (81 year old, AG09908) and a healthy female donor (82 year old, AG08269) were obtained from the Coriell Institute for medical research and grown in DMEM (Sigma-Aldrich, D5671) supplemented with 15% FCS, 2 mM L-glutamine, 100 U/ml penicillin and 0.1 mg/ml streptomycin.

Cryopreserved embryonic cortical neurons isolated from day-17 mouse embryos were purchased from ThermoFisher Scientific (A15586). Cells were revived in Neurobasal Plus Medium (ThermoFisher Scientific, A3582901), complemented with 200 mM GlutaMAX I Supplement (ThermoFisher Scientific, 35050) and 2% B-27 Plus Supplement (ThermoFisher Scientific, A3582801), following manufacturer’s instructions. Neurons were then plated on Poly-D-Lysine (Sigma-Aldrich, P6407)-coated 18mm coverslips at the density of 300,000 neurons/coverslip. Half conditioned medium was changed with fresh medium every third day to feed neurons and prevent excessive evaporation. At 6-days-in-vitro (6 DIV), neurons were transduced with lentiviral particles for the expression of ER-mit RspA-splitFAST (Vector Builder, see plasmids) in conditioned medium supplemented with polybrene (1 µg/ml). 24 hours later, medium was replaced with 0.5 ml of conditioned medium (collected before transduction) and 0.5 ml of fresh medium. Imaging was performed 7 days after transduction.

For astrocyte and microglia primary cultures, we used a previously published protocol^44^, with few modifications. The brains from P0-P2 wild-type (WT) and B6.152H (AD) mice were extracted using forceps and submerged in 30 ml cold dissection medium (DMEM high glucose/pyruvate (Thermo Fisher Scientific, #41966-029) and 0.5% Penicillin-Streptomycin (10,000 U/ml; Gibco). Thereafter, the meningeal layers were completely removed, cortexes were isolated and washed in dissection medium for 2 min to remove blood cells. The cortex was minced and dissociated using a P1000 pipette. The suspension was centrifuged (200 x g, 10 min, RT) and the pellet resuspended in pre-heated 2 ml DMEM high glucose/pyruvate supplemented with 10% heat inactivated FCS, and 0.5% Penicillin-Streptomycin, referred to as supplemented DMEM. Cell clumps were eliminated from the mixed glial culture by passing the suspension through a 70 µm-cell strainer (Corning, #431751), and the suspension was seeded into a Poly-D-lysine (PDL; Sigma-Aldrich, A-003-E)-pre-coated 75 cm^2-^ cell culture flask containing 10 ml of the supplemented DMEM. On day 2, mixed cultures were washed twice with pre-heated D-PBS (pH 7.5; Sigma-Aldrich, D8537) and supplemented DMEM was added. The medium was replaced every 3-4 days.

After 10 days, fresh medium supplemented with 100 ng/mL recombinant murine M-CSF (Peprotech, #315-02), which promotes development and proliferation of microglia^45^, was added. After 4–5 days, microglia was harvested by tapping and seeded to flasks/well plates for further experiments in supplemented DMEM. Microglia harvest was performed up to 4 times in intervals of 2 to 5 days. After final microglia isolation, the remaining attached cells, expected to be enriched in astrocytes, were harvested by 2.5% trypsin/0.5 mM EDTA for 7-10 min at 37 °C and seeded in flasks/well plates for further experiments in supplemented DMEM. Both microglia and astrocytes were transduced with lentiviral particles for expression of ER-mit RspA-splitFAST (Vector Builder, see plasmids) in medium containing polybrene (1 µg/ml). 24 h later, the medium was replaced with fresh medium. Imaging was performed 72 h after transduction. Where indicated, astrocytes were exposed for 16 h to a conditioned medium obtained from either WT CHO cells or 7PA2-CHO cells (containing high levels of naturally generated Aβ peptides), as described^21^.

Animals were bred in-house in individually ventilated cages (IVCs) under specific-pathogen-free (SPF) conditions. Water and food were provided *ad libitum*. All animals were maintained in compliance with the Italian animal protection law and the local animal welfare committee. Organ harvesting was approved by the institutional review.

### Cloning, plasmids, siRNA

Unless differently specified, the plasmids generated in this study have been obtained either by PCR amplification or by gene synthesis (Life Technologies), followed by digestion with specific restriction enzymes and insertion into pcDNA3, allowing mammalian expression under the control of the CMV promoter.

#### ER-NFAST

The ER-membrane targeting sequence from ER-ABKAR (Addgene #61508) was amplified by PCR. The NFAST sequence was amplified by PCR from the Addgene plasmid pAG148 FRB-NFAST (Addgene #130812) and ligated at the 3’ of ER-targeting sequence into pcDNA3.

#### OMM-short-CFAST10

The OMM targeting sequence encoding the N-terminal 72 aminoacids of human TOM70 was amplified by PCR from the previously described TOM70-YFP plasmid^46^, including the CFAST10-encoding sequence in the reverse primer. The fragment was ligated into pcDNA3.

#### OMM-long-CFAST10

The OMM targeting sequence coding the N-terminal 111 aminoacids of human TOM70 was amplified by PCR from the previously described TOM70-YFP plasmid^46^, including the CFAST10-encoding sequence in the reverse primer. The fragment was ligated into pcDNA3. The cytosolic region of TOM70 from the end of the N-terminal transmembrane domain till aminoacid 111 is flexible and works as a linker, as predicted by AlphaFold.

#### *ER-NFAST in pcDNA™FRT⁄TO vector* (used to generate inducible HeLa T-Rex clones)

The sequence encoding ER-NFAST was obtained by enzymatic digestion (HindIII + XhoI) of the above described *ER-NFAST* plasmid and ligated into *pcDNA™FRT⁄TO* vector (ThermoFisher Scientific).

#### ER-frNFAST

The sequence encoding frNFAST was amplified by PCR from the plasmid pAG499-FRB-N-frFAST^13^ and ligated in place of the NFAST sequence in the above described ER-NFAST plasmid.

#### ER-RspA-NFAST

The sequence encoding RspA-NFAST was amplified by PCR from the plasmid pAG573 FRB-RspA(N)-IRES-mTurquoise2^14^ and ligated in place of the NFAST sequence in the above described ER-NFAST plasmid.

#### OMM-short-RspA-CFAST

The OMM targeting sequence coding the N-terminal 72 aminoacids of human TOM70 was amplified by PCR from the previously described TOM70-YFP plasmid^46^, including the RspA-CFAST-encoding sequence^14^ in the reverse primer. The fragment was ligated into pcDNA3.

#### PM-RspA-CFAST

PM-RspA-CFAST was generated by gene synthesis (Life technologies), fusing the RspA-CFAST encoding sequence^14^ at the 3’ of the Lyn11 PM-targeting sequence. The coding fragment was ligated into pcDNA3.

#### MyrPalm-D1cpv

The PM-targeting sequence MyrPalm was amplified by PCR and ligated at the 5’ of the cameleon D1cpv-encoding sequence (Addgene #37479) into pcDNA3.

#### PM-frNFAST

The sequence encoding frNFAST was amplified by PCR from the plasmid pAG499-FRB-N-frFAST^13^ and ligated in place of the RspA-CFAST sequence in the above described PM-RspA-CFAST plasmid. This construct was used to simultaneously detect ER-PM and ER-mit MCSs in the very same cells. For this approach, we used ER-RspA-CFAST (see below) to compliment both PM-frNFAST and OMM-short-RspA-NFAST (see below).

#### ER-RspA-CFAST

ER-RspA-CFAST was generated by gene synthesis (Life Technologies), fusing the sequence encoding RspA-CFAST (ref) at the 3’ of the ER-membrane targeting sequence of ER-NFAST. The insert was then ligated into pcDNA3.

#### OMM-short-RspA-NFAST

The sequence encoding RspA-NFAST was extracted by enzymatic digestion (BamHI + XhoI) from the plasmid ER-RspA-NFAST and ligated in place of the RspA-CFAST-encoding fragment into the OMM-short-RspA-CFAST plasmid described above.

#### OMM-Cepia3

The sequence encoding Cepia3 was amplified by PCR from the plasmid pCMV CEPIA3mt (Addgene #58219) and ligated at the 3’ of the OMM-targeting sequence encoding the N-terminal 72 aminoacids of human TOM70.

#### ER-mit PRINCESS

ER-mit PRINCESS is based on the co-expression of ER-M13-NFAST and OMM-CaM*-CFAST10. To generate ER-M13-NFAST, the M13-encoding sequence was amplified by PCR from the plasmid pCMV CEPIA3mt (Addgene #58219) and ligated between the ER-membrane targeting sequence and the NFAST-encoding sequence in the above described ER-NFAST plasmid. OMM-CaM*-CFAST10 was generated by amplifying by PCR the sequence encoding mutated Calmodulin (CaM*) from the plasmid pCMV CEPIA3mt (Addgene #58219). The CaM*-encoding fragment was then ligated between the OMM-targeting sequence and the CFAST10-encoding sequence in the OMM-short-CFAST10 plasmid described above.

#### ER-PM PRINCESS

ER-PM PRINCESS is based on the co-expression of ER-M13-NFAST (see above ER-mit PRINCESS) and PM-CaM*-CFAST10. To generate PM-CaM*-CFAST10, the CaM*-CFAST10-encoding fragment was extracted by enzymatic digestion from the OMM-CaM*-CFAST10 plasmid (see above ER-mit PRINCESS) and ligated in place of the RspA-CFAST sequence in the above described PM-RspA-CFAST plasmid.

#### OMM PRINCESS

OMM PRINCESS is based on the co-expression of OMM-M13-NFAST and OMM-CaM*-CFAST10 (see above ER-mit PRINCESS). OMM-M13-NFAST was generated by extracting the M13-NFAST-encoding sequence from the plasmid ER-M13-NFAST (see above ER-mit PRINCESS) and ligating it in place of the CFAST10-encoding sequence in the above described OMM-long-CFAST10 plasmid.

#### STIM1-RspA-NFAST

The sequences encoding STIM1 and RspA-NFAST were amplified by PCR from respectively the human STIM1-YFP plasmid (Addgene #19754, a gift from A. Rao) and the pAG573 FRB-RspA(N)-IRES-mTurquoise2^14^ plasmid. The RspA-NFAST-encoding fragment was then ligated at the 3’ of the STIM1-encoding fragment into pcDNA3.

#### STIM1-mit PRINCESS

STIM1-mit PRINCESS is based on the co-expression of STIM1-M13-NFAST and OMM-CaM*-CFAST10 (see above ER-mit PRINCESS). STIM1-M13-NFAST was generated by extracting the M13-NFAST-encoding sequence from the plasmid ER-M13-NFAST (see above ER-mit PRINCESS) and ligating it in place of the RspA-NFAST-encoding fragment in the STIM1-RspA-NFAST plasmid described above.

#### Lentiviral plasmid VB220320-1120bts for the expression of ER-mit RspA-splitFAST

The lentiviral vector VB220320-1120bts was generated by gene synthesis (VectorBuilder) and used to generate lentiviral particles for the expression and/or stable integration of inserts allowing the expression of ER-mit RspA-splitFAST. In this vector, the above described ER-RspA-NFAST and OMM-RspA-CFAST sequences were inserted under the control of respectively CMV and SV40 promoters, allowing the simultaneous expression of the two complimentary fragments. A puromycin-resistance gene was included under the control of mPGK promoter, allowing the selection of cell clones with stable integration of the lentiviral vector.

#### C. elegans plasmid for ER-mit RspA-splitFAST

Plasmid p L4665 (a gift from Andrew Fire, Addgene #1667) carrying the *C. elegans myo 3* promoter was used as backbone to clone an insert allowing expression of the ER-mit RspA-splitFAST. The sequence TOMM70::RspACFAST::SL2::EMB8::RspANFAST was generated by gene synthesis and cloned into pL4665 using BamHI and NcoI restriction enzymes. We used the targeting sequences of *C. elegans* TOMM70 and EMB8 to respectively target RspA-CFAST and RspA-NFAST to the OMM and the ER-membrane. The *C. elegans*-specific SL2 sequence was inserted between TOMM70-RspA-CFAST and EMB8-RspA-NFAST to guarantee co-expression of the two fragments by a single plasmid. The resulting plasmid [Pmyo-3::TOMM70::RspACFAST::SL2::EMB8:: RspANFAST::unc-54 3’UTR, pRF4] allows the expression of ER-mit RspA-splitFAST on body wall muscle cells. Plasmid pRF4 (*rol-6; su1006*) was used as co-injection marker for pHX6743 *C. elegans* strain generation (see below).

The plasmids encoding mit-RFP and H2B-GCaMP6f were previously described^46^. The plasmid encoding OMM-RFP was a gift of G. Hajnoczky. The plasmid encoding ER-GFP (GFP-Sec61β) was a gift of G. Voeltz. The following plasmids were from Addgene: ER-StayGold (Addgene #186296), mCherry-Drp1 (Addgene #49152), Tubuline-GFP (Addgene #56450), LifeActGFP (Addgene #58470).

For RNAi experiments, we used the following siRNA (all from Merck):

MISSION® siRNA Universal Negative Control #1 (SIC001); STIM1 (SASI_Hs01_00107803); STIM2 (SASI_Hs02_00354130); TMEM110 (SASI_Hs01_00204728); SERCA-2A (SASI_Hs01_00047711); ESYT1 (SASI_Hs01_00188940); VPS13A (SASI_Hs02_00306926); VPS13D (SASI_Hs01_00155099).

### Reagents and fluorogens

HMBR (^TF^Lime, 480541-250), HBR-3,5DOM (^TF^Coral 516600-250) and HPAR-3OM (^TF^Poppy 555670-250) were from Twinkle Factory. All the other reagents were from Merck, unless differently specified.

### Fluorescence microscopy

Imaging of the different splitFAST-based probes for MCSs by fluorescence microscopy was performed on a Thunder Imager 3D Cell Culture (Leica), equipped with a LED8 illumination system, both 40x/1.30 oil immersion (HC PL Fluotar 340) and 63x/1.40 oil immersion objective and with a cool camera (Hamamatsu Flash 4.0 V3). Cells seeded and transfected on 18 mm coverslips, were mounted just before the imaging experiments on a chamber and bathed in mKRB (in mM: 140 NaCl, 2.8 KCl, 2 MgCl_2_, 1 CaCl_2_, 10 HEPES, 10 glucose, pH 7.4 at 37°C). Where indicated, CaCl_2_ concentration was 100 µM, or replaced with EGTA 0.5 mM. The probes were complemented with either Lime (4 µM, Twinkle Factory 480541-250) or Coral (4 µM, Twinkle Factory 516600-250) and imaged using respectively the 475 nm (15%) or 555 nm (20%) LED8 excitation lines. Excitation and emission light was filtered by GFP- or TRITC-dedicated sets of filters (Chroma), respectively. In most experiments, a 60 ms exposure time and a frame rate of 15 s/frame were set, unless differently specified (see figure legends). For fast acquisitions, frame rates of 100 ms/frame and 250 ms/frame were used for PRINCESS- or OMM-Cepia3-based experiments, respectively. Where indicated, different cell stimuli/treatments were acutely added to the experimental medium, including histamine (Hist, 100 µM), Thapsigargin (Tg, 100 nM), N,N,N′,N′-tetrakis(2-pyridinylmethyl)-1,2-ethanediamine (TPEN, 500 µM), bradykinin (BK, 100 nM), ATP (100 µM). In some experiments, a 30 min pre-incubation with BAPTA-AM (10 µM), added to mKRB supplemented with CaCl_2_ 1 mM, was performed, as indicated. For the calculation of the genuine change of fluorescence of the different MCS splitFAST-based probes, we performed parallel experiments for each experimental condition/stimulus in cells expressing either cytosolic RspA-NFAST (plasmid pAG573 FRB-RspA(N)-IRES-mTurquoise2)^14^ and RspA-CFAST (plasmid pAG580 FKBP-RspA(C)-IRES-iRFP670)^14^, or NFAST (pAG148 FRB-NFAST, Addgene #130812) and CFAST10 (pAG241 FKBP-CFAST10, Addgene #130814), keeping the same illumination/exposure parameters. The average fluorescent signals of the cells expressing cytosolic RspA-splitFAST (or splitFAST) were subtracted from those of the cells expressing the corresponding MCS probes, to normalize for bleaching and other possible fluorescence changes not linked to MCS remodelling. All the data shown in this manuscript were normalized for the corresponding fluorescence changes of cytosolic RspA-splitFAST (or splitFAST).

### Confocal, STED and FLIM microscopy

Confocal analysis was performed on a Leica TCS-II SP5 STED CW microscope, equipped with a 100x/1.4 N.A. Plan Apochromat objective, a tunable WLL laser, both PMT and HyD (Leica) detectors for signal collection, and a suitable resonant scanner (8,000 Hz) for fast acquisitions. The WLL was set at 488, 555 and 600 nm to respectively excite the different splitFAST-based probes complemented with either Lime (Twinkle Factory 480541-250), Coral (Twinkle Factory 516600-250) or Poppy (Twinkle Factory 555670-250, this latter to complement fr-splitFAST). When a quantitative comparison between different experimental conditions was performed, images were collected by keeping the same excitation/emission, zoom, and image size parameters. Where indicated, 3D-stacks were collected with *z*-steps of 0.4 µm. Images were background-subtracted and analyzed with specific *ImageJ* (NIH) plugins. In particular, the calculation of the percentage of mitochondrial surface in contact with either ER or the PM was performed by staining mitochondria for 15 min in mKRB (or medium) supplemented with MitoTracker Deep Red (300 nM, Thermo Fisher M22426). PM staining was performed incubating the cells in 2 µg/ml WGA (Thermo Fisher W32464) for 25 min in mKRB. After washing, cells were imaged in mKRB, supplemented with Lime (3 µM). Images for both channels were background subtracted and preprocessed using a custom-written Python script. The following dedicated Python libraries have been employed: opencv and scikit-image.

Images were min-max normalized and filtered with a Gaussian filter with a standard deviation of 1 using the “filters.gaussian” function from scikit-image. Images were binarized using a threshold set at half of the mean value of the pixel intensity values exceeding 20% of the pixel intensity distribution.

To locate spatial overlaps between ER-mit or ER-PM RspA-splitFAST and MitoTracker, we merged the binarized images using pixel multiplication. The fraction of overlap has been calculated on the MitoTracker image and the result saved using the ‘to_excel’ function from pandas.

To improve the signal-to-noise ratio, after background subtraction some of the displayed images were further processed with the automatized *ImageJ* plugin *enhance image*, but all analyses were performed before this modification. For fast acquisitions (such as those for the experiments with PRINCESS), resonant scanner was used, with the following parameters: image size 1024×512 pixels, bidirectional acquisition, line accumulation 3x, frame average 3x, 293 ms/frame.

STED and FLIM analyses were performed on a Leica Stellaris 8 TauSTED microscope, equipped with a 100x/1.4 N.A. Plan Apochromat objective (for STED imaging) or a 63x/1.4 N.A. Plan Apochromat objective (for FLIM analysis), a WLL, HyD detectors, a continuous 660 nm STED laser and a pulsed 775 nm STED laser.

For STED analysis of endogenous STIM1 recruitment close to mitochondria, after the indicated treatments to deplete ER Ca^2+^ content (see Fig. 6e legend), HeLa cells were washed with PBS, fixed in 4% formaldehyde (10 min), washed 3x with PBS and permeabilized for 2 h in blocking solution (0.3% Triton X-100, 1% BSA, 5% goat serum in PBS). Cells were incubated overnight at 4°C with primary antibodies (α-TOM20 Santa Cruz # sc-11415; α-STIM1 Merck WH0006786M1) diluted (1:200) in antibody dilution buffer (0.3% Triton X-100, 1% BSA in PBS), washed 3 x 5 min with PBS and incubated for 1 h at RT with secondary antibodies (α-mouse STAR 580 and α-rabbit STAR 635P, Abberior) diluted in antibody dilution buffer. Coverslips were washed 4 x 5 min in PBS and mounted with Aqua-Poly/mount (Polysciences, 18606). STED images were acquired by sequential acquisition of each channel, setting the WLL at respectively 570 nm and 635 nm, by activating (30%) the 775 nm pulsed depletion-laser. Excitation/emission parameters were kept constant, avoiding signal saturation. Pixel size was set at 20 nm/pixel. Images were background-subtracted, and the Manders’ and Pearson’s co-localization coefficients calculated by the *ImageJ Co-localization Analysis* plugin.

For FLIM analysis, we used the Phasor Method^47^, available within the FALCON>FLIM application module of the Leica Application Suite X (4.6.0) software. Briefly, each pixel of the FLIM image was transformed into a unique position in the phase diagram by Fourier transformation of the decay curve, with the x-axis representing the real component and the y-axis the imaginary component. The different lifetime components were visualized as clustering of pixels in defined regions of the phasor plot, that were isolated by adjusting the size and position of specific cursors. The different lifetime components, identified by each of these cursors, were then broken down into separate channels by the “Separate” function provided by the FALCON module. To ascertain the molecular identity of each specific component, control experiments were performed by preparing samples in which single splitFAST-based probes were expressed separately.

### Lattice Light-sheet and Airyscan microscopy

HeLa cells stable clones were seeded on µ-Dish 35 mm (Ibidi) and 24-48 h later incubated for 10 minutes in culture media (37 °C, 5% CO_2_) with MitoTracker Deep Red (200 nM, Thermo Fisher M22426). Imaging was performed at room temperature in mKRB using ZeissLattice Lightsheet 7, to get gentle volumetric live imaging. ER-mit RspA-splitFAST (marked with 3 µM ^TF^Lime) and MitoTracker Deep Red signals were simultaneously acquired in two channels using a dual camera configuration with bandpass 505-545 nm, and 656-750 nm detection. Excitation was combined 488 nm and 640 nm with a Sinc3 30×1000 lightsheet. Excitation intensities were balanced to minimize photobleaching and maintain a good signal to noise ratio. Integration time was set to 15 ms. The imaged volumes were recorded in 500 slices with step 0.2 µm and pixel size 0.145 µm, Time lapse recordings were made with 6 volumes per minute for 10 minutes. Data were postprocessed by iterative deconvolution followed by deskewing, resulting in a coverslip transformed image volume of 297×100×38 µm for each time point Surface rendering was performed by IMARIS software after median (3×3×3) filtering of the ER-Mit RspA-splitFAST signal to reduce background noise. Surface tracking was calculated using the “Connected Components” algorithm and data collected from IMARIS “Statistics” module.

COS-7 cells were seeded on µ-Dish 35 mm (Ibidi), co-transfected 24 h later with plasmids encoding ER-Mit RspA-splitFAST and either ER-StayGold, mCherry-Drp1, Tubulin-GFP or LifeActGFP. The day after, cells were incubated for 10 minutes in culture media (37 °C, 5% CO2) with MitoTracker Deep Red and then imaged at room temperature in mKRB (supplemented with either 3 µM Lime or 3 µM Coral) using a Plan-Apochromat 63×/1.4 NA oil objective on an inverted Zeiss 980 LSM Airyscan2 confocal microscope. Multi-color timelapse imaging was performed by sequentially scanning three channels configured for 639 nm excitation with longpass 660 nm detection, 561 nm excitation with shortpass 615 nm detection, and 488 nm excitation with bandpass 495-555 nm detection. For 3D timelapse imaging, a full z-stack of 5-15 slices was acquired per laser line before switching. Zoom factor was set at 5x yielding maximum pixel-dwell time of ∼0.69 μs per pixel and a frame time 325 ms at Nyquist sampling (∼40 x 40 x 150 nm per voxel). High resolution images were obtained by Airyscan processing in Zen Blue (3.8) using standard 3D Auto defined strength. Surface rendering was performed by using IMARIS software.

To segment ER structures of COS-7 cells into “sheets” (rough ER) and “tubules” (smooth ER), we have used a size-based approach. On average, the tubules measure between 10-12 pixels in our recordings (Resolution: 23.4989 pixels per micron), while the sheets are bigger. The range between the biggest and the smallest tubule structure is 5 pixels to 21 pixels. On average the tubules range a width of 10.451 ± 2.5172 pixels, mean ± SEM, n=5 cells. Using the erode function in ImageJ, we created a mask to erode 6 times (eliminating structures <12 pixels) to only keep the sheets. The sheets are then removed from the original ER channel to only keep the tubules. In this manner we can separate the different structures from each other to make co-localization analysis with the mitochondria and split-fast signal in the other channels. Structures <5 pixels were removed and regarded background. To analyze the pictures in a systematic way an ImageJ macro was constructed and applied to all the films. The full macro can be found at: https://github.com/linneapavenius/Segmentation-of-ER-sheets-and-tubules-.

After quantification, performed as detailed above, representative images and videos were obtained calibrating the signal through the IMARIS “Display Adjustment” console.

The Kernel Density and the correlation Spearman index were calculated by Excel XLSTAT (Microsoft).

### Flow cytometry

14 days after selection with either Hygromycin (300 µg/ml, HeLa T-Rex clones expressing ER-mit splitFAST; induction of ER-mit splitFAST expression by overnight treatment with tetracyclin 2 µg/ml) or puromycin (3 µg/ml, for stable HeLa and COS7 cell clones expressing ER-mit RspA-splitFAST after transduction with lentiviral particles, see plasmids), cells were bathed in PBS supplemented with Lime (5 µM). The cells expressing the probe of interest were separated by Fluorescence-activated cell sorting (FACS), using a S3e TM Cell Sorter (BIORAD) endowed with a 488 nm laser and a 525/30 nm emission filter. Analysis was performed with ProSort 1.6 (BIORAD). After sorting, single cell clones were isolated from the positive cell population by limiting dilution cloning in 96 well-plates, followed by a screening for optimal probe-expression performed by fluorescence microscopy.

### CLEM

HeLa cells were seeded on a 35mm Dish with gridded glass coverslip (MatTek In Vitro Life Science Laboratories) and transfected with plasmids to express ER-mit RspA-splitFAST. 24 h after transfection, cells were incubated for 20 min with MitoTracker™ Red CMXRos (200 nM, Thermo Fisher M7512) to stain mitochondria, washed in PBS and bathed in PBS containing Lime (5 µM). Confocal images of cells expressing ER-mit RspA-splitFAST were acquired by a FluoVIEW 3000 RS system (Evident Scientific, Waltham, MA) equipped with a UPLSAPO 60X/1.3 Silicon objective. The entire thickness of the cell was acquired with optical section depth of 0.21 μm and the images were deconvolved with Huygens Software before alignment with the matching EM images (SVI, Hilvertum, Netherlands). After the acquisition of the full z-stack, fixative was added (final concentration 4% PFA in 0.1 M Hepes) and the field was acquired for further 10 minutes. Dishes were removed from the microscope stage and postfixed with 2,5% glutaraldehyde in 0.1 M cacodylate buffer (pH 7.4) for 1 h at RT. Samples were postfixed with reduced osmium (1% OsO4, 1.5% potassium ferrocyanide in 0.1 M cacodylate buffer, pH 7.4) for 2 h on ice. After several washes in milli-Q water, sections were incubated in 0.5% uranyl acetate overnight at 4°C. Samples were then dehydrated with increasing concentration of ethanol, embedded in epoxy resin and polymerized for 48 h at 60 °C. Ultrathin serial sections were obtained using an ultramicrotome (UC7, Leica microsystem), collected on formvar-carbon coated copper slot grids, stained with uranyl acetate and Sato’s lead solutions and observed in a Transmission Electron Microscope Talos L120C (FEI, Thermo Fisher Scientific) operating at 120kV. Images were acquired with a Ceta CCD camera (FEI, Thermo Fisher Scientific) and aligned with confocal images using ec-CLEM plugin of ICY software. To evaluate the fraction of mitochondrial surface in contact with ER, analysis was performed with Microscope Image Browser (MIB) software^48^.

### Generation and maintenance of *C. elegans* strains

*C. elegans* strains were cultured and maintained following standard methods as previously described^49^. The **pmyo3::**ER-mit RspA-splitFAST allele *sybIs6743* (see in plasmids) was generated at SunyBiotech (http://www.sunybiotech.com) by plasmid microinjection together with a *rol6* co-injection marker, followed by X-ray irradiation to randomly integrate the sequence in the worms genome and generate a stable *C. elegans* strain. All experiments were performed on synchronized worms generated by L4 larvae isolation, and maintained at 20°C.

*C. elegans* worms (N2, control not injected; pHX6743, *sybIs6743*) were evaluated in a 3% agarose pad slide, pre-treated with Levamisole 1 mM, and surrounded by a thin silicone line. When levamisole was completely absorbed by the agarose pad, 3 μL of the fluorogen solution (^TF^Coral 40 μM in M9 Buffer) were placed in the center of the pad and the worms transferred into the solution. The worms were covered with a cover slip and incubated O/N on the fluorogen solution. Imaging was performed 16 h later.

### Ca^2+^ measurements

For Aequorin Ca^2+^ measurements, HeLa cells (0.6×10^5^ cells/well) were plated on coverslips (13 mm diameter) and transfected with plasmids for the expression of either cytosolic Aequorin or mitochondrial mutated Aequorin, together with the indicated siRNA (see above).

48 h after transfection, cells were incubated for 1 h at 37°C in mKRB supplemented with native coelenterazine (5 µM, BIOTIUM), and then incubated for 15 minutes in mKRB supplemented with either TPEN (500 µM, to induce partial depletion of ER Ca^2+^ content) or EtOH (as control). After incubation with EtOH, control cells were additionally incubated for 30 s in mKRB supplemented with TPEN (500 µM). This brief treatment was performed to match the possible contribution of residual TPEN in buffering Ca^2+^ peaks, while not being long enough to induce an appreciable increase of ER-Mit juxtaposition, which was observed to only occur at least 4-5 minutes after depletion of ER Ca^2+^. The coverslips were then transferred to the perfusion chamber and perfused with mKRB. Histamine (100 µM) was added in Ca^2+^-free, EGTA-containing mKRB. The experiments ended by permeabilizing cells with digitonin (100 µM) in a hypotonic Ca^2+^-rich solution (10 mM CaCl_2_ in H_2_O) to discharge the remaining aequorin pool and calibrate the signal. Luminescence was collected by a photon counter (Package Photon Counting 9125, Sens-Tech) and analysis performed by the associated software (Sens-Tech). The light signal was analysed and converted into Ca^2+^ concentrations as described^50^.

ER Ca^2+^ content was measured by ratiometric GEM-CEPIA1er^32^. HeLa cells, expressing GEM-CEPIA1er and treated as indicated, were mounted into an open-topped chamber in mKRB and imaged by a Thunder Imager 3D Cell Culture (Leica), equipped with a LED8 illumination system, a 40x/1.30 oil immersion (HC PL Fluotar 340) objective and a cool camera (Hamamatsu Flash 4.0 V3). Excitation was performed by the 390 nm LED8 line, and emissions at either 530 or 480 nm were simultaneously collected by a beam splitter (502 nm dichroic mirror, 535/30 nm and 480/40 nm emission filters, Chroma Technologies). Exposure time was 60 ms; images were acquired every 2.5 seconds. After background subtraction, the ratio between the blue and the green emission (proportional to ER Ca^2+^ concentration) was calculated by the LASX software (Leica).

For ratiometric OMM-Cepia3 measurements, HeLa cells expressing OMM-Cepia3 and ER-mit RspA-splitFAST were mounted into an open-topped chamber in mKRB supplemented with ^TF^Coral (5 µM) and imaged by a Thunder Imager 3D Cell Culture (Leica), equipped with a LED8 illumination system, a 63×/1.40 oil immersion (HC PL Fluotar 340) objective and a cool camera (Hamamatsu Flash 4.0 V3). ER-mit RspA-splitFAST was excited with the 555 nm led line and emission collected by a TRITC filter (Chroma Technologies), whereas OMM-Cepia3 was sequentially excited with the 475 nm (10 ms exposure) and 390 nm (50 ms exposure) led lines (in both cases, we used a 495 nm dichroic mirror and a 525/50 nm emission filter). Indeed, we exploited the isosbestic point of Cepia3 (∼395 nm) to perform ratiometric Ca^2+^ measurements, sequentially exciting at 475 nm (Ca^2+^-sensitive wavelenght) and at 390 nm (Ca^2+^-insensitive wavelenght). The overall frame rate was 100 ms/frame. A 2×2 binning was set. Images were background substracted and the ratio between the two wavelenghts performed by the LASX software. To specifically evaluate the OMM-Cepia3 ratio at the level of ER-mit MCSs, we used the “AND” function of *ImageJ* between the OMM-Cepia3 ratio image and the corresponding ER-mit RspA-splitFAST image.

### Statistical analysis

All data are representative of at least 3 independent experiments. When a pairwise comparison with the control group was performed, significance was calculated by unpaired Student’s *t*-test for normally distributed and Wilcoxon Mann–Whitney U test for not normally distributed data. When more than two groups were compared, statistical significance was calculated using two-way analysis of variance (ANOVA) with post-hoc comparisons (Tukey), setting p<0.05 as a level of significance. Statistical analysis was performed by GraphPad Prism software v9.2. * = p < 0.05, ** = p < 0.01, *** = p < 0.001. Values are reported as mean ± SEM.

## Acknowledgements

This study is dedicated to the memory of Tullio Pozzan, a great scientist, a mentor and a dear friend for many of us. The authors gratefully acknowledge L. Ozmen and F. Hoffmann-La Roche Ltd (Basel, Switzerland) for kindly donating the AD mouse model used in this study (MTA 16-02-07-UniPD), E. Trevisson and V. Morbidoni (University of Padua) for helping with the setting of the *C. elegans* strains, P. Romani for helping with the setting of confocal microscopy, D. Sandonà for suggestions on cloning strategies and P. Magalhães for reading the manuscript. Part of the schemes presented in this work were created with BioRender.com.

P.G.C fellowship was supported by Ayudas de recualificación del Sistema Universitario Margarita Salas (2021-2023), University of Valladolid-Spanish Ministry of Education, financed by the European Union through EU Next Generation. This work was supported by grants from the Italian Ministry of University and Scientific Research (PRIN2017XA5J5N); the University of Padova (SID2019); Cure Alzheimer’s Fund (USA) to P.P.; the Dynamic Imaging program of the Chan-Zuckerberg Initiative DAF (grant number 2023-321185), an advised fund of Silicon Valley Community Foundation, to A.G. and R.F; the Italian Ministry of University and Scientific Research grant (PRIN P20225R4Y5, financed by the European Union, NextGenerationEU) to R.F. The authors acknowledge Euro-BioImaging (www.eurobioimaging.eu) for providing access to imaging technologies and services via the *Advanced Light Microscopy Italian Node* (*Laboratory of Ca^2+^ and cAMP signaling in physiology and pathology*, Padua, Italy; and *ALEMBIC*, Milan, Italy); the Swedish National Microscopy Infrastructure, NMI (VR-RFI 2019-00217); EuroBioImaging-ERIC and the PNRR infrastructure, SEELIFE n. IR00023 (financed by the European Union, NextGenerationEU).

## Author contributions

P.G.C., M.R., P.P., and R.F. were responsible for experimental design and data interpretation. P.G.C., M.R., L.P., M.S., H.B. and R.F. performed and analyzed experiments. N.A and S.S prepared primary glial and neuronal cell cultures. M.B. conceived the script for analyzing images. V.B. and A.R. performed CLEM experiments and image analysis. M.S. performed AlphaFold analysis. M.L.S., P.A., B.A.N. provided reagents, suggestions and discussed experiments. L.N. and M.A. provided suggestions and discussed experiments. A.G. provided all the original splitFAST-based tools, reagents, expertise on splitFAST and suggestions. P.P. and R.F. conceived the research and secured funds. R.F. wrote the manuscript with the help of the other authors. All the authors have agreed to publish the current version of the manuscript.

## Competing interests

A.G. is co-founder and holds equity in Twinkle Bioscience/The Twinkle Factory, a company commercializing the FAST and split-FAST technologies. The remaining authors declare no competing interests.

