## Supplemental Material for "Simultaneous detection of membrane contact dynamics and associated Ca^2+^ signals by reversible chemogenetic reporters"

### Supplementary Figures, Table and Supplementary Video legends

#### Supplementary Fig. 1

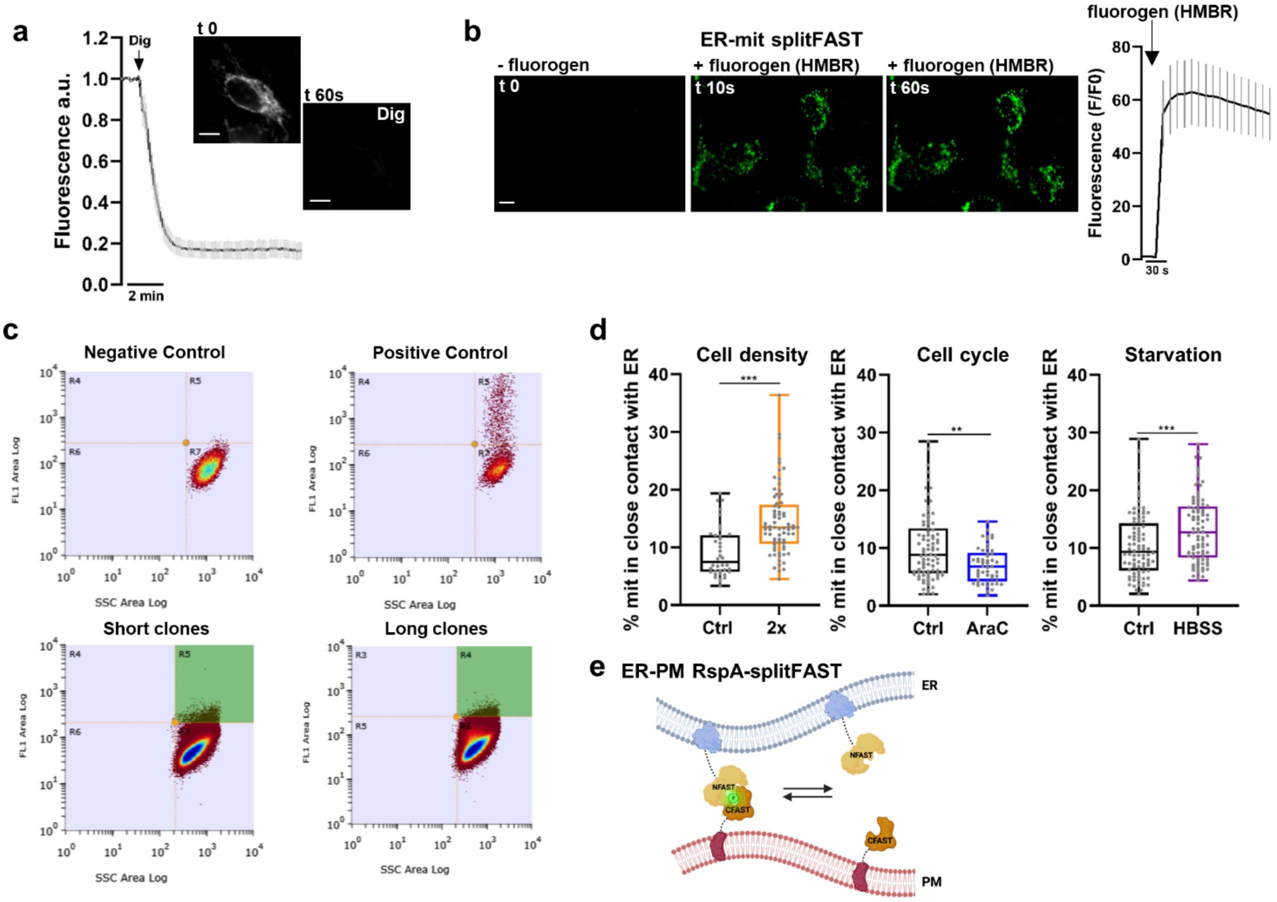


**Supplementary Figure 1: The splitFAST system is a versatile and reversible system to study MCSs. a,** Representative images and mean ± SEM trace of the fluorescent signal in HeLa cells, expressing the CFAST10 fragment targeted to the OMM and the NFAST fragment free in the cytosol. In this condition, the mitochondrial network is marked by reconstituted splitFAST. Few seconds after cell permeabilization by digitonin (25 µM), the release of the cytosolic NFAST fragment in the extracellular *milieu* induces a fast drop in the fluorescent signal, confirming that NFAST-CFAST10 assembly is fully and promptly reversible. n=12 cells from 3 independent experiments. Scale bar, 10 µm. See also the corresponding Supplementary Video 1. **b,** Representative images and mean ± SEM trace of the fluorescent signal in HeLa cells expressing ER-mit splitFAST, before and upon addition of the fluorogen HMBR (3 µM, see Methods), as indicated. n=21 cells from 3 independent experiments. Scale bar, 10 µm. **c,** FACS analysis (side scatter SSC-A versus fluorescence FL1-A) of HeLa T-Rex cells (see Methods), either not transfected (negative control), transiently transfected with ER-mit splitFAST (positive control) or selected with antibiotics after transfection with either the short or long ER-mit splitFAST. In all the conditions, tetracycline (16 h) was used to induce expression of the different splitFAST probes. Cells were bathed in PBS supplemented with the fluorogen HMRB (5 µM). The signal of ER-mit splitFAST (green area on the bottom panels) was used to sort the cells expressing either short or long ER-mit splitFAST. Single clones were then isolated by limiting dilution. **d,** Box plots represent the percentage of mitochondrial surface engaged in the formation of close contacts with ER, as observed in stable, inducible HeLa cell clones expressing short ER-mit splitFAST and treated as indicated (see Methods), in which mitochondria were marked by MitoTracker Deep Red. Higher cell density (2x cell density, corresponding to 90% confluency), as well as starvation (HBSS, 40 min, 37°C), increase ER-mit MCSs, whereas cell cycle blockage by

AraC (3µM, 16 h) decreases this parameter. n>38 cells from at least 3 different experiments **$p$< 0.01; ***$p$< 0.001. **e,** The cartoon represents the tailoring of the RspA-splitFAST system to mark ER-PM MCSs.

#### Supplementary Fig. 2

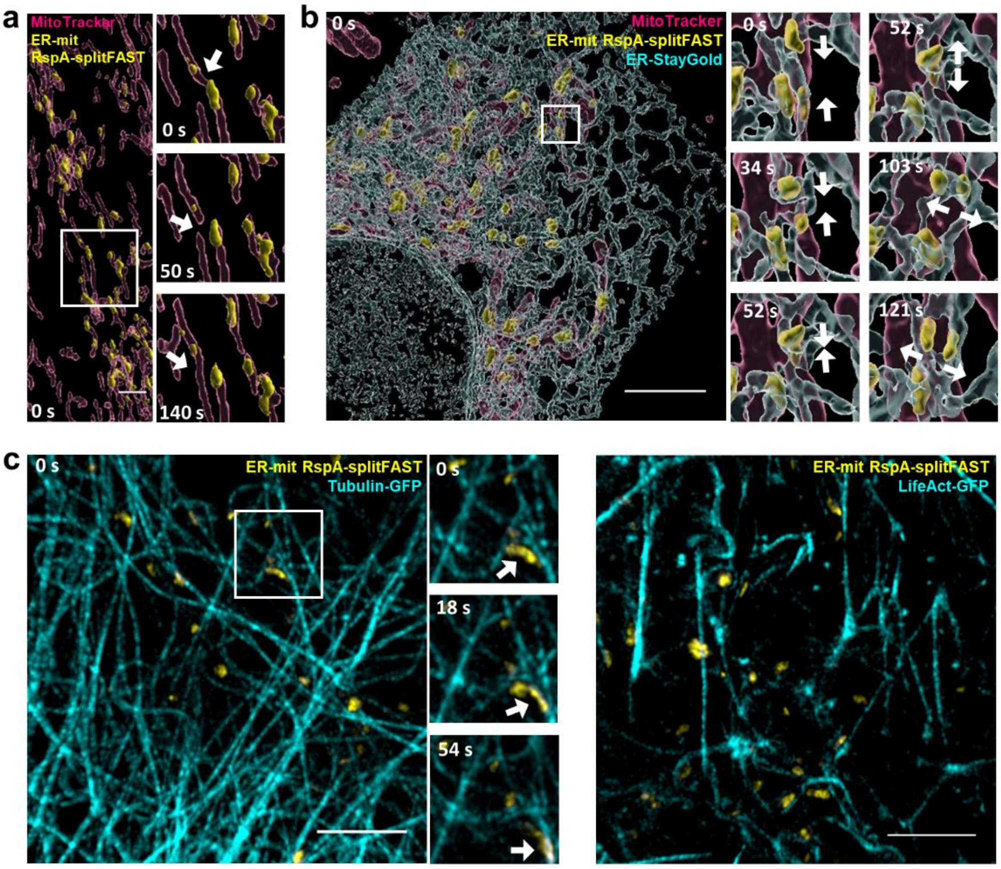


**Supplementary Figure 2: ER-mit MCS dynamics and association with cytoskeleton. a,** Representative lattice light-sheet 3D image of a region of a HeLa cell, expressing ER-mit RspA-splitFAST (surface rendering, yellow) and stained with MitoTracker Deep Red (surface rendering, magenta). Enlarged regions show the time-lapse of an ER-mit MCS (arrow) at a mitochondrial fission site (0-50 s) and another MCS (arrow) at a mitochondrial fusion site (50-140 s). **b,** Representative Airyscan 3D image of a COS-7 cell expressing ER-mit RspA-splitFAST (surface rendering, yellow), ER-StayGold (surface rendering, cyan) and stained with MitoTracker Deep Red (surface rendering, magenta). Enlarged regions show the time-lapse of two ER-mit MCSs (arrows) fusing together upon fusion of two ER tubules (0-52 s), following by a fission of the very same MCS (arrows) upon ER tubule fission (52-121 s). See also the corresponding Supplementary Video 9. **c,** Representative Airyscan 3D images of a COS-7 cell expressing ER-mit RspA-splitFAST (yellow), Tubulin-GFP (cyan, left image) or LifeAct-GFP (cyan, right image). ER-mit MCSs locate in proximity of microtubules and actin filaments. Enlarged regions show the time-lapse of an ER-mit MCS moving along a microtubule (arrow). Scale bar, 5µm. See also the corresponding Supplementary Video 10.

### Supplementary Fig. 3

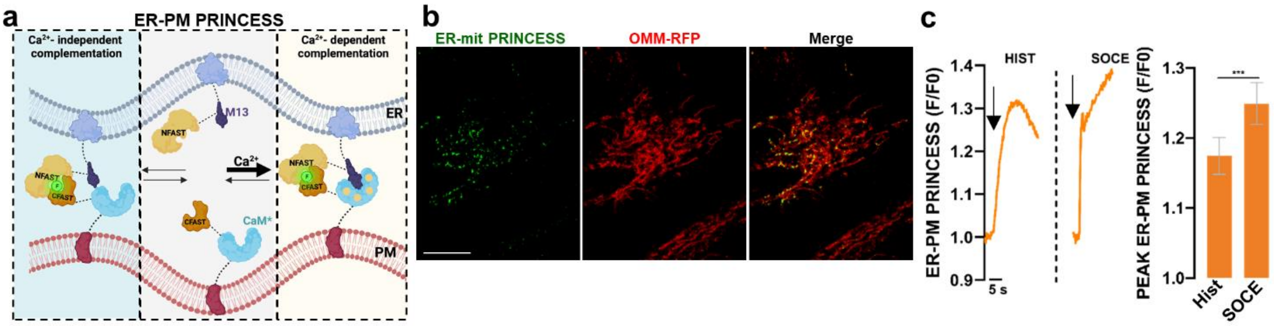


**Supplementary Figure 3: ER-PM PRINCESS to measure $Ca^{2+}$ dynamics at ER-PM MCSs. a,** The cartoon represents the rationale behind the design of ER-PM PRINCESS (see also Fig. 5c legend). **b,** Representative confocal images of ER-mit PRINCESS fluorescent signal in HeLa cells, co-expressing ER-mit PRINCESS and OMM-RFP. **c,** Representative traces of ER-PM PRINCESS fluorescence increases in HeLa cells, upon histamine (100 µM) stimulation in $Ca^{2+}$-free mKRB (see Methods), or $CaCl_2$ (2 mM) addition (SOCE) after 6 min depletion of ER-$Ca^{2+}$ content (obtained by histamine (100 µM) and thapsigargin (100 nM) stimulation in $Ca^{2+}$-free mKRB). On the right, bars represent the peaks of ER-PM PRINCESS fluorescence (expressed as F/F0) upon the indicated treatments. Mean ± SEM; n>36 cells from 3 independent experiments. ***p<0.001.

### Supplementary Fig. 4

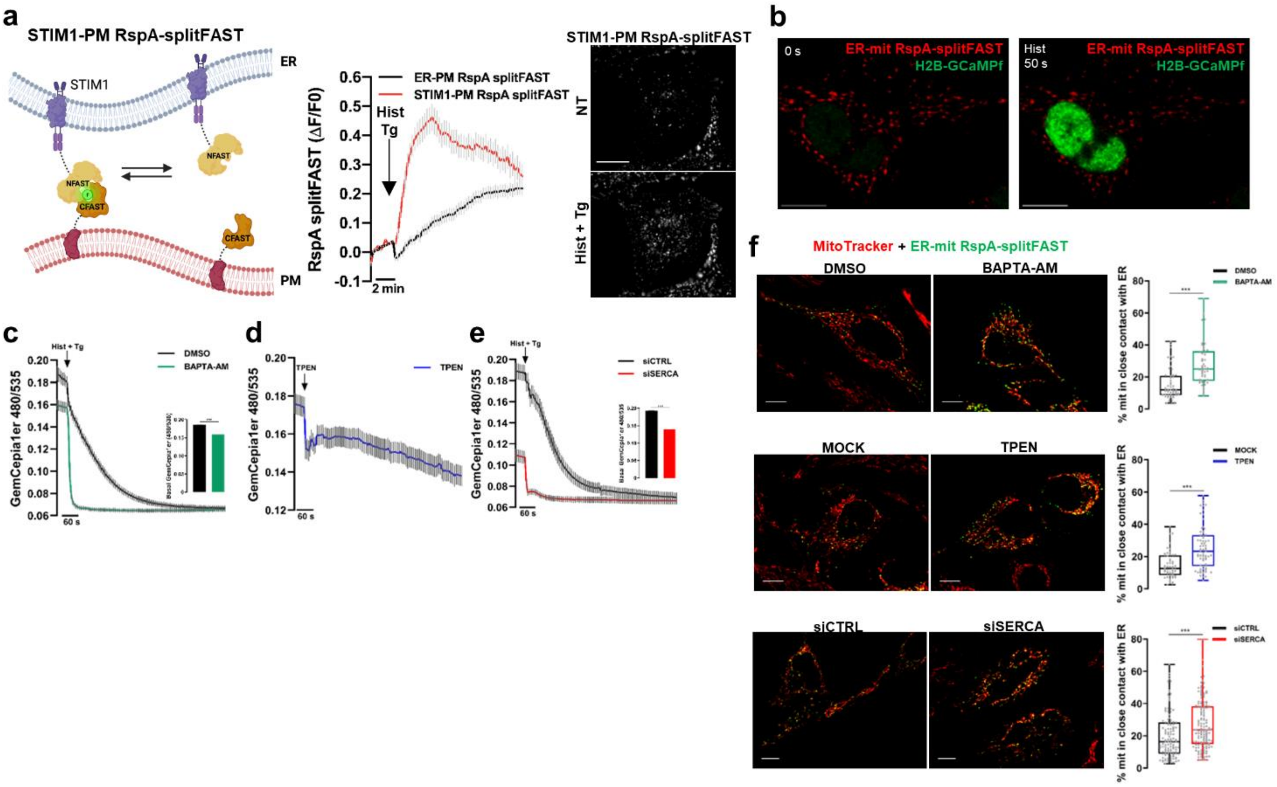


**Supplementary Figure S4: ER $Ca^{2+}$ depletion induces ER-mit MCS remodelling. a**, **d**, On the left, the cartoon represents the tailoring of the RspA-splitFAST system to detect the recruitment of STIM1 close to PM. In the middle panel, mean ± SEM traces of the fluorescent signal in HeLa cells expressing either the ER-PM or the STIM1-PM RspA-splitFAST probe, upon ER $Ca^{2+}$ depletion (100 µM histamine plus 100 nM thapsigargin, arrow). n>55 cells from 3 independent experiments. On the right, representative images of HeLa cells expressing STIM1-PM RspA-splitFAST, before (NT, not treated) or 180 after stimulation with histamine plus Tg (Hist + Tg). **b**, Representative confocal images of HeLa cells co-expressing the ER-mit RspA-splitFAST probe (red) and the nuclear $Ca^{2+}$ sensor H2B-GCAMP6f (green, to detect cytosolic $Ca^{2+}$ elevations, which are in equilibrium with the nuclear ones). Histamine (100 μM) was added to trigger cytosolic/nuclear $Ca^{2+}$ peak. Scale bar, 10 μm. See also the corresponding Supplementary Video 14. **c-e**, Mean ± SEM traces (and relative basal values in the histograms) of GemCepia1er 480/535 nm ratio emission (proportional to ER $Ca^{2+}$ levels, see Methods) in HeLa cells stimulated with 100 μM histamine plus 100 nM thapsigargin (Hist + Tg). Cells were treated with the vehicle (DMSO, black trace in **c**), or BAPTA-AM (10 μM for 30 min before measurements, green trace in **c**), or 500 μM TPEN (blue trace in **d**), or transfected with control- (black trace in **e**) or SERCA2A-specific siRNA (red trace in **e**). In **c, e,** the bars on the right represent the basal GemCepia1er 480/535 nm ratio for the indicated conditions, before stimulation with Hist + Tg. Mean ± SEM; n>63 cells in **c**, >62 cells in **d**, >23 cells in **e**, from 3 independent experiments. **f**, Representative confocal images of HeLa cells, expressing the ER-mit RspA-splitFAST probe and labelled with MitoTracker Deep Red, in the indicated conditions (as in **c-e**). Scale bar, 10 μm. The box plots represent the percentage of mitochondrial surface (% mit) engaged in close contact with the ER in each condition. Mean ± SEM; n>53 cells (DMSO vs BAPTA-AM), >52 cells (MOCK vs TPEN), >118 cells (siCTRL vs siSERCA), from 3 independent experiments. ***$p<0.001$.

### Supplementary Fig. 5

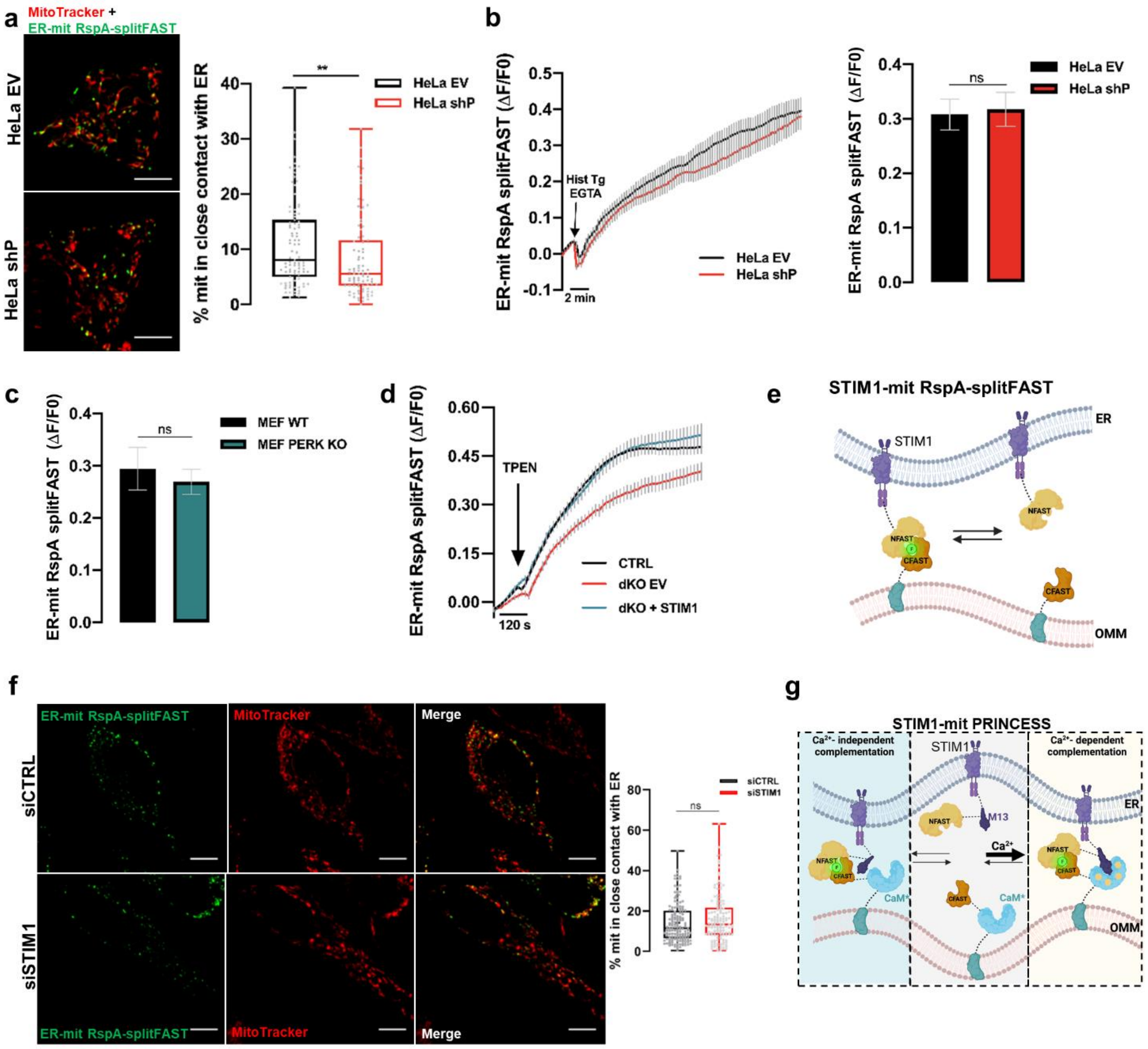


**Supplementary Figure 5: $Ca^{2+}$-regulated remodelling of ER-mit MCSs is mediated by STIM1**
**a,** Representative confocal images of HeLa cell clones, either control (EV) or with stable PERK-downregulation (shP), expressing ER-mit RspA-splitFAST and in which mitochondria were stained by MitoTracker Deep Red. On the right, the box plots represent the percentage of mitochondrial surface co-localized with ER-mit RspA-splitFAST for the indicated cell types. n>87 cells from 3 independent experiments. **p < 0.01. **b,** Mean ± SEM traces of ER-mit RspA-splitFAST fluorescent signal in EV or shP HeLa cell clones, upon depletion of ER $Ca^{2+}$ content by histamine (Hist, 100 µM) and thapsigargin (Tg, 100 nM) stimulation in a $Ca^{2+}$-free, EGTA-containing mKRB. On the right, bars represent the change, for the indicated cell types, of ER-mit RspA-splitFAST fluorescent signal (expressed as ΔF/F0) 15 min after Hist + Tg stimulation. Mean ± SEM; n>88 cells from 3 independent experiments. **c,** Bars represent the change, for the indicated cell types, of ER-mit RspA-splitFAST fluorescent signal (expressed as ΔF/F0) 15 min after ATP (100 µM) + Tg (100 nM) stimulation. Mean ± SEM; n>25 cells from 3 independent experiments. **d,** Mean ± SEM traces of ER-mit RspA-splitFAST fluorescent signal in SH-SY5Y cell clones control (CTRL) or dKO for STIM1/STIM2, in which STIM1 was re-expressed (dKO + STIM1) or not (dKO EV), upon acute depletion of ER $Ca^{2+}$ content by exposure to TPEN (500 µM). n>44 cells from 3 independent experiments. **e,** The cartoon represents the rationale behind the design of a RspA-splitFAST-based

probe to monitor STIM1 recruitment close to mitochondria. RspA-NFAST was fused at the C-terminus of WT STIM1 and can complement with OMM-RspA-CFAST when close enough. **f,** Representative confocal images of HeLa cells, co-transfected with the cDNAs for the expression of the ER-mit RspA-splitFAST and either control siRNA (siCTRL) or STIM1-specific siRNA (siSTIM1) and in which mitochondria were marked with MitoTracker Deep Red. On the right, box plots represent the percentage of mitochondrial surface co-localized with ER-mit RspA-splitFAST. Mean ± SEM; n>139 cells from 3 independent experiments. **g,** The cartoon represents the rationale behind the design of STIM1-mit PRINCESS (see also Fig. 5c legend).

**Supplementary Table 1**

| Downregulated Protein | n | ER-mit RspA-splitFAST ΔF/F0 | p value | Ref |
|---|---|---|---|---|
| CTRL | 24 | 0.215 ± 0.0326 | - | - |
| SERCA-2A | 35 | 0.213 ± 0.0366 | 0.8723 | 1 |
| STIM2 | 22 | 0.213 ± 0.0445 | 0.6047 | 2 |
| TMEM110 | 26 | 0.104 ± 0.0357 | *0.0318 | 3 |
| STIM1 | 35 | 0.0948 ± 0.0309 | **0.0074 | 2 |
| ESYT1 | 41 | 0.196 ± 0.0411 | 0.7937 | 4 |
| VPS13A | 30 | 0.183 ± 0.0335 | 0.5449 | 5 |
| VPS13D | 27 | 0.238 ± 0.0411 | 0.6234 | 6 |

| Cell clone | n | ER-mit RspA-splitFAST ΔF/F0 | p value | Ref |
|---|---|---|---|---|
| HeLa EV | 88 | 0.3079 ± 0.0282 | - | 7 |
| HeLa shP | 95 | 0.3175 ± 0.0312 | 0.8723 | 7 |
| MEF WT | 25 | 0.2944 ± 0.0408 | - | 7 |
| MEF PERK KD | 34 | 0.2693 ± 0.02405 | 0.9333 | 7 |

**Supplementary Table 1:** The table shows the results of a siRNA-based screening for possible molecular candidates involved in the $Ca^{2+}$-dependent modulation of ER-mit MCSs. The results are expressed as change of ER-mit RspA-splitFAST fluorescence (ΔF/F0) after 15 min from stimulation of ER-$Ca^{2+}$ depletion, either by histamine + Tg (HeLa cells) or ATP + Tg (MEF cells). Mean ± SEM; displayed n correspond to the number of cells, from at least 3 independent experiments.

#### Supplementary Video legends

**Supplementary Video 1:** The Video corresponds to the experiment described in Supplementary Fig. 1a.
**Supplementary Video 2:** The Video corresponds to the experiment described in Fig. 2b.
**Supplementary Video 3:** The Video corresponds to the experiment described in Fig. 3b.
**Supplementary Video 4:** The Video corresponds to the experiment described in Fig. 3c.
**Supplementary Video 5:** The Video corresponds to the experiment described in Fig. 3d.
**Supplementary Video 6:** The Video corresponds to the experiment described in Fig. 3e.
**Supplementary Video 7:** The Video corresponds to the experiment described in Fig. 3f.
**Supplementary Video 8:** The Video corresponds to the experiment described in Fig. 3g.
**Supplementary Video 9:** The Video corresponds to the experiment described in Supplementary Fig. 2b.
**Supplementary Video 10:** The Video corresponds to the experiment described in Supplementary Fig. 2c.

**Supplementary Video 11:** The Video corresponds to the experiment described in Fig. 4d.
**Supplementary Video 12:** The Video corresponds to the experiment described in Fig. 5e.
**Supplementary Video 13:** The Video corresponds to the experiment described in Fig. 5e
**Supplementary Video 14:** The Video corresponds to the experiment described in Supplementary Fig. 4b.
